# Charting the Multi-level Molecular Response to Palbociclib in ER-Positive Breast Cancer

**DOI:** 10.1101/2025.08.27.672587

**Authors:** Archishma Kavalipati, Amy Aponte, Michael E. Sullivan, Sarah L. Whittington, José C. Martínez, Grant Goda, Maria M. Aleman, Michael J. Emanuele, Daniel Dominguez

## Abstract

The addition of CDK4/6 inhibitors to endocrine therapy has significantly improved outcomes in HR+/HER2-breast cancer. However, variable patient responses and acquired resistance remain a clinical challenge. We therefore defined the comprehensive molecular response to palbociclib, the most clinically used CDK4/6 inhibitor. Global analyses of gene expression, protein abundance, splicing, and chromatin accessibility revealed broad patterns and specific changes that result from CDK4/6-inhibition in breast cancer cells. We uncovered unexpected feedback between CDK4/6 and estrogen-response signaling, which has clear clinical implications. We also revealed a widespread alternative splicing program that partially overlapped with genes whose expression is regulated, and which is expected to impact protein function. These molecular changes nominated combination therapies that interfere with the activation of CDKs or ERα. Accordingly, co-targeting CDK7, which regulates CDK2, CDK4/6 and ERα, additively impacted cell fitness. Collectively, these data reveal a complex, multi-tiered response to CDK4/6 inhibition, with implications for therapeutic efficacy.

## INTRODUCTION

Breast cancer (BC) is the second most prevalent cancer and second leading cause of cancer deaths among women in the U.S.^1^. This highly heterogenous disease can be divided into subtypes characterized by expression of the hormone receptors (HR), estrogen receptor (ER), progesterone receptor (PR), as well as the receptor tyrosine kinase, human epidermal growth factor (HER2/ERBB2). Patients with HR+/HER2-disease account for approximately 70% of breast cancer cases. In HR+ BC, activation of the ER or PR pathways stimulates production of D-type cyclins. Cyclins D1, D2 and D3 bind to and activate the cyclin-dependent kinases 4 and 6 (CDK4/6), and this complex phosphorylates and inactivates the retinoblastoma (RB) tumor suppressor, leading to its dissociation from the E2F family of transcription factors and promoting transcription of genes necessary for DNA replication and cellular proliferation [extensively reviewed in ^2–5^].

The cyclin D-CDK4/6-RB axis is therefore an attractive target for cancer therapy, and the FDA has approved three CDK4/6 inhibitors (CDK4/6i): palbociclib (hereafter Palbo), ribociclib and abemaciclib^6^. These inhibitors prevent the phosphorylation of RB and result in cell cycle arrest, thereby suppressing tumor growth. Hormone therapy in combination with CDK4/6 inhibition is the primary first-line treatment for HR+/HER2-advanced or metastatic BC^7^. Clinical data demonstrates that in combination with endocrine therapies, CDK4/6i significantly increases progression-free and overall survival in patients^8,9^. Despite these improvements in patient outcomes, like other targeted therapies, CDK4/6i present various clinical challenges, including variable patient responses and resistance, both innate and acquired. Notably, up to 35% of breast cancer patients treated with CDK4/6i never respond, and acquired resistance is common among nearly all patients^10,11^. Resistance to CDK4/6i is associated with various molecular alterations, including loss of RB protein, cyclin E1 overexpression, and amplifications of *CCND1*, *CDK4, and CDK6* [reviewed in ^12^]. Nevertheless, there are no approved molecular biomarkers to select patients for treatment, beyond subtyping as HR+/HER2-. Thus, while the cyclin D-CDK4/6-RB axis and its relationship to hormone receptor signaling has been well documented, there remain unknown aspects of this pathway which influence the effectiveness of these therapies in patients.

A complete cataloging of the molecular responses to CDK4/6i at the transcriptional, post-transcriptional, and proteomic level is critical in understanding the mechanisms that underlie response and resistance to CDK4/6i. In this study, we present a multi-omics analysis on HR+/HER2-BC cells, focusing on changes in gene expression, alternative splicing, and proteomics. Our analysis revealed pathways and networks altered in response to CDK4/6 inhibition, and we find evidence for a negative feedback loop between CDK4/6 and ER. This suggests that hyper-activated hormone-dependent signaling might occur in patients in response to CDK4/6i and, in turn, influence the response to pathway inhibition and possibly resistance. This dynamic feedback is reminiscent of the activation of insulin signaling through the PI3K-mTOR pathway in response to PI3K inhibition^13^. Motivated by findings in our multi-omics analysis, we find that co-inhibition of CDK4/6 and CDK7 using the selective inhibitor and clinical candidate samuraciclib additively suppresses proliferation. Collectively, our findings reveal that CDK4/6i remodel the transcriptomic and proteomic landscape in ER+ BC cells, and we provide evidence that supports further evaluation of potential co-targets to improve clinical outcomes.

## RESULTS

### Cell cycle-related mRNA transcripts are downregulated upon palbociclib treatment

To measure transcriptional and proteomic changes after Palbo, we treated the well-established ER+/HER2-T-47D BC cell line with DMSO (control) or 1 µM Palbo in biological triplicate. After 24 hours, we collected samples and performed stranded paired-end RNA sequencing and whole-cell mass spectrometry on samples prepared in parallel (**Fig. 1A**). To understand the transcriptome-wide response to Palbo treatment, we performed differential expression analysis using DESeq2^14^. PCA data visualization showed clear separation between treatment groups (**Fig. S1A**, upper), and after filtering for well-detected transcripts, we detected 19,460 genes. Of these, 1,056 were found to be differentially expressed (Table S1). Furthermore, most differentially expressed genes were downregulated versus upregulated (833 down vs. 233 up), a statistically significant direction (p = 2.79e-83, exact binomial test).

**Figure 1:**
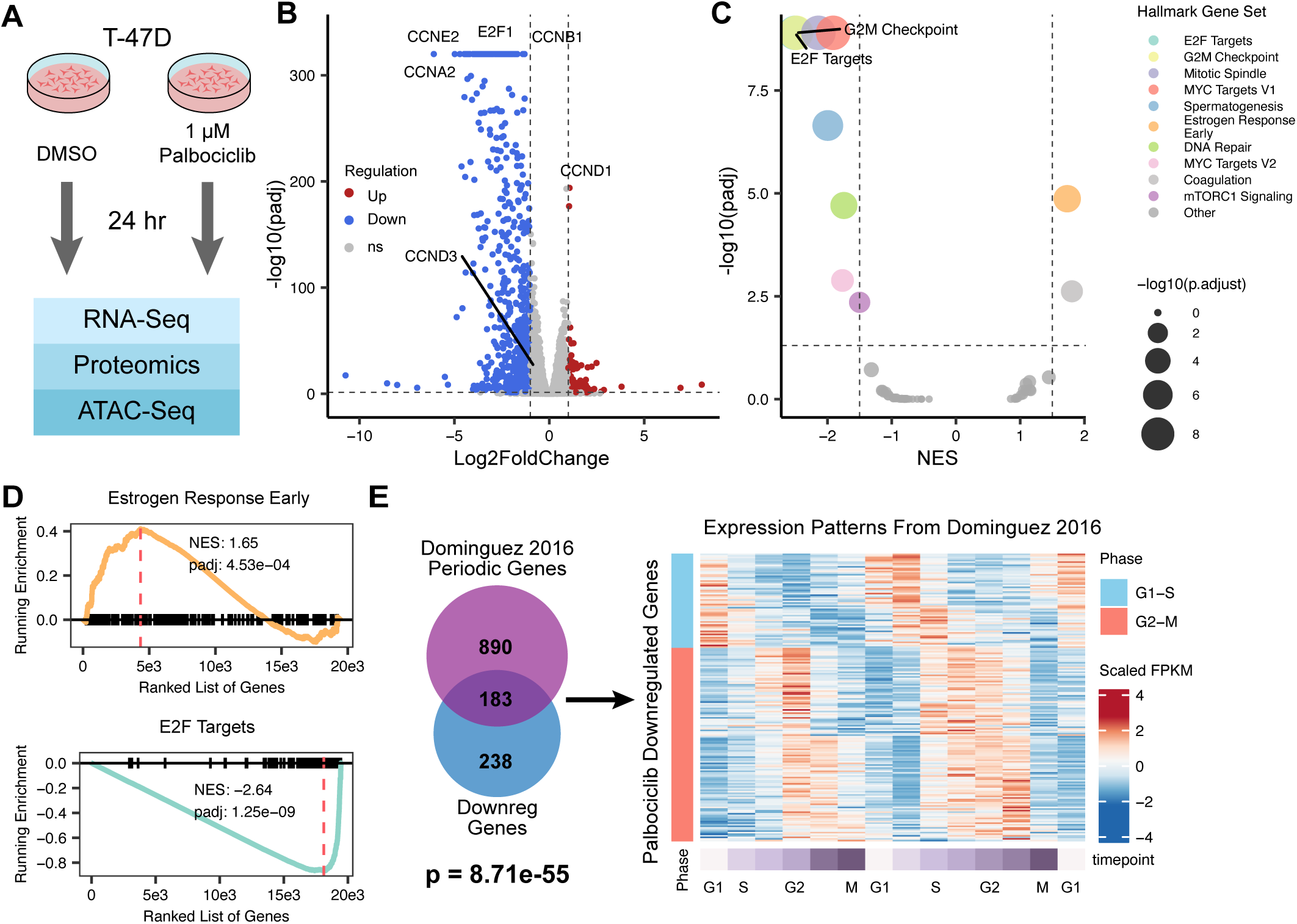
Cell cycle-related mRNA transcripts are downregulated upon Palbo addition. **A.** Experimental schematic. HR+/HER2-T-47D breast cancer cells were treated with DMSO (control) or 1μM Palbo for 24 hours before sample collection and subsequent RNA-seq, ATAC-seq, and proteomics analysis. **B.** Volcano plot showing differential expression of genes after Palbo treatment. Blue points represent downregulated genes (Down); red points represent upregulated genes (Up); grey points are not significant (ns). **C.** Plot showing GSEA of all genes after Palbo treatment using hallmark gene sets. Genes were ranked by rank = log2FoldChange * (1 - p-value). A normalized enrichment score (NES) ≥ abs(1.5) and a p-adjusted value ≤ 0.05 were used as the cutoff values for a gene set to be labeled. **D.** Leading edge plots of two hallmark gene sets shown in 1C. Upregulated genes are enriched for “Estrogen Response Early” while downregulated genes are enriched for “E2F Targets.” **E.** Left: Overlap between periodically expressed genes identified by Dominguez et al. 2016 and genes downregulated after Palbo treatment (p = 1.36e-44, hypergeometric test). Right: Row-scaled gene expression values (FPKMs) of these 183 genes across 2 cell divisions.

Consistent with the well-known role of CDK4/6 in cell cycle control and in transcriptional regulation of S-phase genes, established cell cycle genes were strongly downregulated after Palbo treatment (**Fig. 1B**). These included the transcriptional activator *E2F1*, and cyclins, including G1-S transition-associated *CCNE1*, S phase-associated *CCNA1*, and the mitotic *CCNB1*. Gene Ontology (GO) analysis of downregulated genes (**Methods**) revealed significant enrichment for cell division-related terms (**Fig. S1B**), including chromosome segregation (GO:0007059, p-adjusted = 6.14e-66), organelle fission (GO:0048285, p-adjusted = 5.14e-57), nuclear division (GO:0000280, p-adjusted = 8.57e-61), and DNA replication (GO:0006260, p-adjusted = 8.97e-51). Surprisingly, despite most G1-S transition genes being downregulated, we found significant upregulation of *CCND1* (**Fig. 1B**), suggestive of a unique regulatory mechanism in this context (discussed in more detail below).

To better understand enriched pathways across the Palbo-responsive transcriptome, we performed Gene Set Enrichment Analysis (GSEA) with a focus on hallmark gene sets from^15^ (**Methods**). We found 10 enriched hallmark gene sets, 8 of which were in downregulated genes and 2 in upregulated genes (**Fig. 1C**, abs(NES) > 1.5 and p-adjusted ≤ 0.05). The most enriched hallmark term in the downregulated gene set was “E2F Targets” (p-adjusted = 1.25e-09, **Fig. 1D** lower), underscoring that the anti-proliferative effect of inhibiting CDK4/6 activity occurs largely through suppression of E2F-mediated transcription. Other enriched hallmark terms among downregulated genes were closely associated with cell cycle progression, including G2/M Checkpoint, Mitotic Spindle, Myc Targets, and DNA Repair.

CDK4/6i predominantly arrest cells in G_1_, so among genes expected to change after CDK4/6 inhibition are those that fluctuate in expression depending on cell cycle stage, hereafter referred to as periodic genes^16^. We sought to understand which of these periodic genes were most impacted by Palbo. Using a previous analysis defining transcriptome-wide changes in periodic gene expression during cell cycle progression^17,18^, we compared periodically expressed genes with Palbo-responsive differentially expressed genes. While Palbo-upregulated genes showed little overlap with periodic expressed genes, our analysis revealed 183 genes that were both Palbo-downregulated and periodic (p = 8.71e-55, hypergeometric test. These genes showed a clear periodic trend in their expression patterns over the cell cycle (**Fig. 1E**). We further stratified the periodic genes based on their peak expression stage, G1-S or G2-M, and computed their overlaps with Palbo-downregulated genes to find 60 and 123 overlaps respectively (**Fig. S1C**). We expanded our periodic analysis to four additional cell cycle-dependent gene expression datasets^18–21^(**Methods**) and corroborated overlaps with Palbo-downregulated genes (**Fig. S1D**). Although relatively few genes were found in common between Palbo-downregulated genes and all surveyed cell cycle datasets, those that did overlap have critical roles in cell division. These include *CENPQ*, *CCNE1*, *CCNF*, *POLD3*, *AURKB*, and *MKI67*, further supporting Palbo downregulates transcripts recurrently identify as contributors to cell cycle progression.

To understand upstream control of gene expression changes through chromatin accessibility, we conducted an ATAC-seq analysis on T-47D cells using the same conditions as our previously described RNA-seq. Once again, we identified a strong separation between treatment groups, indicating a clear Palbo-induced effect (**Fig. S1A**, lower). After quantifying accessibility of detected chromatin regions with and without Palbo treatment (Table S2), we observed a similar number of regions that tended toward higher accessibility as we did lower accessibility. Overlapping these chromatin regions with genomic regions from RNA-seq (**Methods**) revealed that certain differentially expressed genes, including some cell cycle genes, overlap differentially accessible genomic regions (**Fig. S1E**). However, we noted only a modest correlation between differentially expressed genes and differentially accessible regions, indicative of limited chromatin accessibility changes during the 24-hour period (**Fig. S1F,** Pearson’s R = 0.26, p < 2.2e-16). Differences in accessibility were generally limited, suggesting that over a 24-hour Palbo treatment period, large changes in chromatin accessibility have yet to be established or detectable by ATAC-seq.

### CDK4/6i triggers an altered splicing program affecting RNA quality control and cell cycle genes

Mechanisms of RNA processing, such as alternative splicing, are known to modulate both mRNA and protein expression but are often overlooked in regulatory studies. To this end, we sought to understand how Palbo impacts post-transcriptional gene regulation through changes in alternative splicing. Using the rMATS software^22^, we identified 1,905 differential splicing events from our RNA-seq (Table S3) across four different event types: alternative 3’ splice sites (A3SS), alternative 5’ splice sites (A5SS), retained introns (RI) and skipped exons (SE). While skipped exons were the predominantly detected event type (1,304 significant events detected overall), we also detected 234 retained intron events (**Fig. 2A**). In fact, a larger fraction of expressed and quantified introns changed in splicing pattern after Palbo treatment than all other splicing event types (*e.g.,* skipped exons), indicating a preferential effect of CDK4/6i on RI as opposed to other events (**Fig. S2A**). This finding is consistent with prior work demonstrating that cell cycle and proliferative genes display regulated intron retention during transitions from resting to proliferative states^23–25^. Next, we sought to determine if alternatively spliced genes were also differentially expressed. Overlap between alternatively spliced genes and differentially expressed mRNAs was minimal (**Fig. 2B**), consistent with previous literature showing that splicing and gene expression responses generally involve non-overlapping genes^26^. However, genes that were impacted through expression and splicing had well-known cell cycle related functions, including components of the minichromosome maintenance complex, mitotic kinases, and centromere proteins, further supported by GO analysis of these overlapping genes (**Fig. 2C**). Thus, the Palbo response involves over a thousand alternative splicing changes, some of which are found in critical cell cycle and proliferation genes.

**Figure 2:**
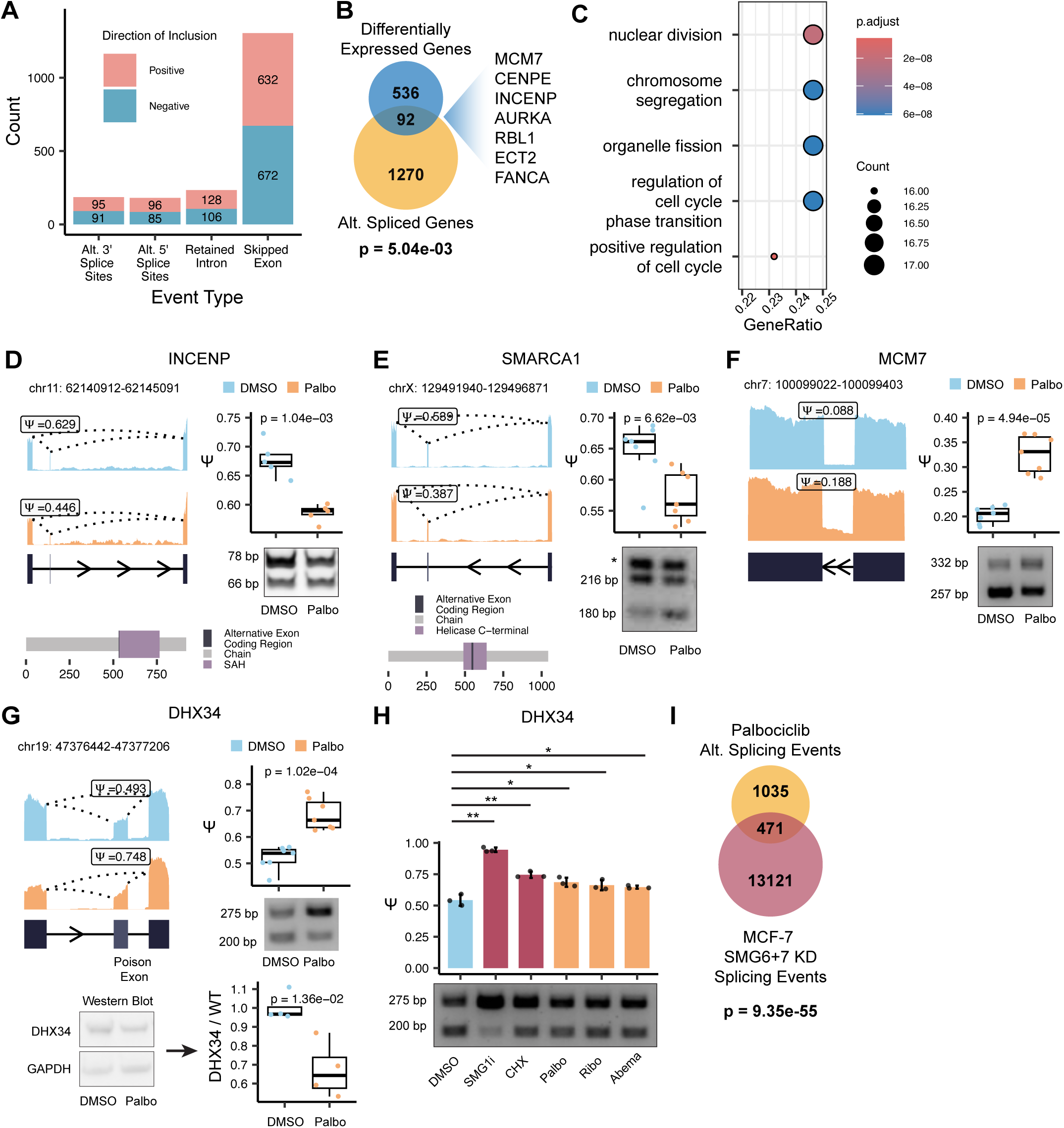
CDK4/6i triggers an altered splicing program affecting cell cycle genes and RNA quality control genes. **A.** Barplot of alternative splicing events detected as significant by rMATS. Positive = more included in Palbo. Negative = less included in Palbo. **B.** Overlap between differentially expressed genes as described in Fig. 1 and alternatively spliced genes. p = 5.04e-03, hypergeometric test. **C.** GO analysis of genes affected at both the expression and alternative splicing level. **D.** Left: visualization of RNA sequencing coverage over INCENP exon skipping event. Right: validation of alternative splicing by RT-PCR and corresponding quantification (p = 1.04e-03, Student’s two-sided t-test). Lower: Alternative exon coding region in INCENP protein. **E.** Left: visualization of genome coverage in SMARCA1 exon skipping event. Right: validation of alternative splicing by RT-PCR and corresponding quantification (p = 1.58e-02, Student’s two-sided t-test). Lower: Alternative exon coding region in SMARCA1 protein. **F.** Left: Visualization of genome coverage in MCM7 retained intron event. Right: validation of alternative splicing by RT-PCR. **G.** Left: visualization of genome coverage in DHX34 exon skipping event. Right: validation of alternative splicing by quantitative PCR. p = 7.5e-05, Student’s two-sided t-test. Lower: validation of lowered DHX34 protein expression by Western blot. p = 1.36e-02, Student’s two-sided t-test. **H.** RT-PCR of DHX34 after treatment with CDK4/6i and NMD inhibitors. All conditions compared to DMSO, p-values calculated via Student’s t-test. SMG1i: p = 1.41e-03, CHX: p = 5.69e-03, Palbo: p = 1.42e-02, Ribo: p = 2.77e-02, Abema: p = 4.89e-02. **I.** Overlap between Palbo-induced splicing events in T-47D and NMDi-induced splicing events in MCF-7. p = 9.35e-55, hypergeometric test.

Previous work has shown that alternative splicing itself can be cell cycle-dependent or periodic^17,20,27^. Accordingly, we explored whether Palbo-responsive splicing events overlap known periodic splicing events. We found that the overlap between periodically spliced (reported in^17^) and Palbo-responsive splicing gene sets was small (**Fig. S2B**, p = 3.34e-04, hypergeometric test). More importantly, the overlap between individual splicing events (*i.e.*, specific exons or introns) in both datasets was both minimal and non-significant (**Fig. S2C**). These data suggest that the splicing response to Palbo treatment is not merely due to altered cell cycle progression; rather, it represents a unique splicing program. A similarly limited overlap was recently reported for splicing events responsive to AURKA inhibition and those that are cell cycle-dependent^27^, supporting a role for cell cycle kinase signaling in the control of unique splicing programs.

Splicing event inclusion (*e.g.,* an exon or intron) is commonly represented by a “percent spliced in” value, denoted by PSI or Ψ, which represents the percentage of all transcripts that include the exon or intron. Relative changes in inclusion level between conditions (e.g. Palbo-treated vs. untreated) can be compared by determining the inclusion level difference (ΔΨ) between conditions. After assessing genes with large ΔΨ between Palbo-treated and DMSO-treated conditions, high RNA-seq read coverage over the splicing event, and functional relevance, we experimentally validated splicing events in 6 candidate genes (**Methods**). Following Palbo treatment, *INCENP*, a component of the chromosomal passenger complex (CPC), showed skipping of exon 11, which encodes a portion of the single α-helix (SAH) domain (**Fig. 2D**, Student’s t-test, p = 1.04e-03). The SAH domain in INCENP functions as a stretchable helix that can that binds microtubules, and its loss could impair microtubule binding of the CPC, as well as the ability of Aurora B to find and phosphorylate substrates^28–30^ Moreover, INCENP is strongly downregulated at the transcriptomic and proteomic levels after Palbo treatment, and it is known that reduction of one component of the CPC can reduce abundance of the other components ^31^. *SMARCA1*, a chromatin remodeler and member of the SWI/SNF family displayed skipping of exon 13 upon Palbo treatment (**Fig. 2E**, Student’s t-test, p = 6.62e-03); the exon skipping isoform has been shown to have chromatin remodeling activity as a functional ATPase, while the exon including isoform loses this function and has oncogenic properties ^32,33^. We identified other events in mitotic genes, such as a Palbo-induced skipping of exon 2 (in the 5’ UTR) of ECT2, a signaling factor involved in cytokinesis^34^ (**Fig. S2D**) and three Palbo-induced exon skipping events in AURKA (**Fig. S2E**). While RT-PCR validation showed Palbo treatment induced exon skipping, individual isoform measurements were not possible due to similarity in sizes; however, skipping was expected for all events based on RNA-seq. Overall, Palbo elicits a broad splicing regulatory program involving cell cycle regulatory factors.

Isoform expression can be controlled at the level of splicing and/or via differential stability/degradation of specific isoforms. For example, the nonsense-mediated decay (NMD) pathway is a surveillance mechanism responsible for degrading isoforms that contain premature termination codon, such as mRNAs containing retained introns, or exons that introduce a premature termination codon (hereafter, poison exons)^35^. We identified and validated increased intron retention after Palbo in *MCM7* (**Fig. 2F**), which encodes an essential protein for genome replication^36^. We also found increased inclusion of a poison exon in *DHX34*, a DEAD-box RNA helicase that is itself involved in the NMD pathway^37^ (**Fig. 2G**, upper). Both intron retention and poison exon inclusion can lead to downregulation of mRNA and protein^38^; indeed, we observed downregulation of MCM7 at the mRNA and proteomic levels (discussed below) and validated downregulation of DHX34 via western blotting (**Fig. 2G**, lower). To verify that DHX34 poison exon inclusion was NMD-sensitive, we treated T-47D cells with an inhibitor of SMG1, a kinase required for the NMD pathway^39,40^. Because NMD is coupled to translation, we additionally treated cells with the translational inhibitor cycloheximide (CHX), a known method to decrease NMD activity. NMD-disrupting treatments led to increased inclusion of the DHX34 poison exon, verifying its NMD sensitivity. Furthermore, treatment with ribociclib or abemaciclib also showed that the inclusion of this exon is affected by multiple CDK4/6 inhibitors (**Fig. 2H**), highlighting the specificity to CDK4/6 inhibition.

Next, we considered whether Palbo-responsive splicing events were generally targeted by the NMD pathway on a global scale. To this end, we obtained RNA-sequencing data from MCF-7 (ER+/HER2-) cells depleted for SMG6 and SMG7^41^, two critical factors for NMD pathway activity^42^. This analysis revealed 471 overlapping Palbo-induced and NMD-regulated alternative splicing events (**Fig. 2I**, p = 9.35e-55, hypergeometric test). Thus, CDK4/6 inhibition alters splicing outcomes of a large set of isoforms that are also sensitive to NMD inhibition suggesting a relationship between cell cycle and quality control pathways.

### CDK4/6 inhibition promotes a global decrease of proliferation-related proteins

Proteome remodeling throughout the cell cycle depends on gene expression, splicing, and dynamic changes in protein stability through targeted, ubiquitin-mediated protein degradation. Thus, changes to the proteome can be controlled independently of changes in gene expression. We performed whole-cell mass spectrometry to determine global proteomic changes in Palbo-treated cells (**Fig. 1A**). In total, we analyzed 7,042 expressed proteins; 345 proteins exhibited significant changes in abundance after Palbo treatment (using p-adjusted value = 0.05 and abs(log2FoldChange) ≥ 0.6), and many more exhibited subtle expression changes (Table S4). In accordance with our transcriptomic expression data, we found a strong and statistically significant downregulation of protein expression in response to Palbo treatment (p = 2.05e-17, exact binomial test, **Fig. 3A**). Among the proteins most strongly downregulated after Palbo treatment, we identified key regulators of cell cycle progression and division.

**Figure 3:**
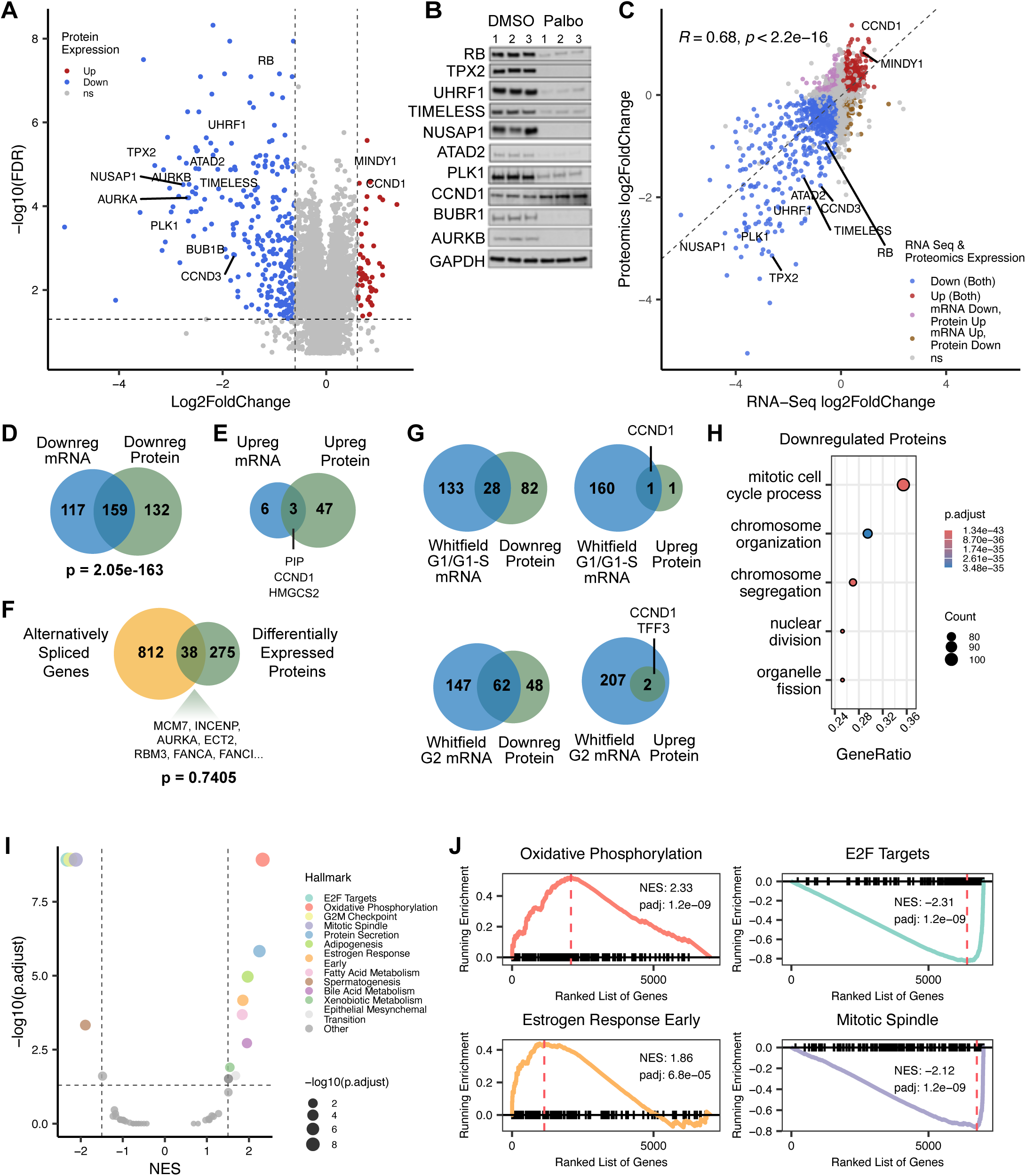
CDK4/6 inhibition promotes a global decrease of proliferation-related proteins. **A.** Volcano plot of differential proteins detected after mass spectrometry (MS). Blue points represent downregulated genes (Down); red points represent upregulated genes (Up); grey points are not significant (ns). **B.** Western blot validation showing expression of indicated proteins from samples used for MS. **C.** Scatter plot showing Palbo-induced gene expression trends at the mRNA (x-axis) and protein (y-axis) levels. Gene directionality was labeled if meeting FDR ≤ 0.05. Pearson’s R = 0.68, p < 2.2e-16. Legend description: Down, negative mRNA and proteomics log2FoldChange. Up, positive mRNA and proteomics log2FoldChange. mRNA down, protein up: negative mRNA log2FoldChange, positive proteomics log2FoldChange. mRNA up, protein down: positive mRNA log2FoldChange, negative proteomics log2FoldChange. ns: not significant. **D.** Overlap between significantly (log2FoldChange ≤ -1, FDR ≤ 0.05) downregulated mRNAs and significantly (log2FoldChange ≤ -0.6, FDR ≤ 0.05) downregulated proteins (p = 2.05-163, hypergeometric test). **E.** Overlaps between significantly upregulated mRNAs (log2FoldChange ≥ 1 and FDR ≤ 0.05) and significantly upregulated proteins (log2FoldChange ≥ 0.6 and FDR ≤ 0.05). **F.** Overlaps between alternatively spliced genes and differentially expressed proteins (p = 0.7405, hypergeometric test). **G.** Overlaps between differentially expressed proteins after Palbo treatment and periodic genes identified by Whitfield et al. 2002. **H.** Top GO terms enriched in downregulated proteins. The p-value and q-value cutoffs were both set to 0.05 and the ontology “BP” (biological process) was used. The top 5 enriched terms are shown. **I.** Plot showing GSEA of all proteins after Palbo treatment using hallmark gene sets. **J.** Leading edge plots of four representative hallmark gene sets. Left: upregulated proteins are enriched for “Oxidative Phosphorylation” and “Estrogen Response Early.” Right: Downregulated proteins are enriched for “E2F Targets” and “Mitotic Spindle”.

Among the most significantly decreased proteins were those that function in chromosome segregation, including components of the CPC (INCENP, AURKB, BIRC5/Survivin, CDCA8/Borealin), inner kinetochore (CENPU, CENPH, CENPN, CENPK), outer kinetochore (NDC80, NUF2, SPC25, SPC25, KNL1, KNL2, MIS18a, MIS18b), the kinetochore-associated fibrous corona (CENPE, CENPF, ZWINT) and components of the mitotic checkpoint (BUB1B/BUBR1 and MAD2L1/MAD2). Also significantly downregulated were key regulators of mitotic spindle function, including but not limited to TPX2, NUSAP1, KIF2C/MCAK, KIFC1/HSET, KIF11/EG5, KIF22/KID, and PRC1. Similarly, there was a significant decrease in the abundance of proteins that carry out DNA replication, among which were TONSL, TIPIN, TIMELESS, PCLAF/PAF15, RFC1, RFC4, POLA2, POLE2, POLE4, GINS1-4, MCM2-7, and PRIM1-2. Moreover, many proteins involved in repair of DNA damage were decreased Palbo treatment and this included BRCA1, FANCA, FANCI, XRCC3, CHK1, LIG1, RMI2, RAD18, and WRN.

Signaling proteins were highly altered in response to Palbo treatment. This included proteins involved in cell cycle phosphorylation: PKMYT1, CCNB1, CCNB2, TTK/MPS1, CKS1B, CKS2, MASTL/GREATWALL, and PLK1. Interestingly, CDK1 and CDK2, which do not oscillate in abundance during a normal cell cycle, were also significantly decreased, consistent with a recent proteomic study in RPE1 cells treated with Palbo^43^. We also observed a decrease in several regulators of ubiquitin signaling, including several E2-ubiquitin conjugating enzymes (UBE2T, UBE2C, UBE2S, UBE2B) as well as E3 ligases or their associated substrate receptors (CDC20, DTL/CDT2, RNF168, RNF4, RNF149).

Finally, we identified several proteins involved in scaffolding of chromosome in mitosis through cohesion or condensation (PTTG1/Securin. NCAPD3, NCAPG2, NCAPH2, SMC4) and others involved in chromatin regulation or dynamics (UHRF1, HELLS, ATAD2, CHAF1A, CHAF1B, MKI67, DNMT1, NSD2, ASF1B, SLBP, SUV39H2/KMT1B, SETMAR).

To validate protein changes from our global mass spectrometry, we probed for candidate proteins by Western blot (**Fig. 3B**). Our validation showed robust changes reflective of our mass spectrometry results, including the increase of cyclin D1 and the decrease of RB. Also downregulated were TIMELESS, a key regulator of DNA replication and the DNA damage response; TPX2 and NUSAP1, regulators of mitotic spindle assembly and dynamics; and the multi-functional mitotic kinase, PLK1. Additionally, we observed the downregulation of UHRF1, a chromatin and DNA methylation regulator which we previously showed is ubiquitinated and degraded in cycling G1-phase cells^44^, and ATAD2, a chromatin remodeling AAA ATPase. The downregulation of these and other chromatin associated proteins in Palbo-treated cells supports studies by our group and others that there are significant changes to the chromatin-regulatory apparatus during cell cycle progression, and in this case, in response to cell cycle arrest^44–46^. This could suggest chromatin readers, writers or erasers as possible co-vulnerabilities in Palbo treated cells.

### Comparison of global mRNA and protein changes

There are various levels of regulation between gene transcription and final protein product. For this reason, we investigated the relationship between transcriptomic and proteomic changes following Palbo treatment. We found a strong correlation between the two datasets (**Fig. 3C**, R = 0.68, p < 2.2e-16), suggesting that Palbo influences protein expression, in part, by regulating RNA expression. We examined the factors that displayed differential abundance only at the transcriptomic or proteomic levels and those that change at both. A subset of genes showed decreased transcriptomic expression but increased proteomic expression (**Fig. S3B**); these genes include *GGH*, a glutamate-hydrolyzing enzyme associated with poor clinical outcomes in invasive breast tumors^47^; *ADIPOR1*, a regulator of glucose and lipid metabolism^48^; *PXMP4*, a peroxisomal gene predicted to be involved in metabolic process^49^ and *HMGN2/3*, a nucleosome binding protein involved in chromatin regulation^50^. Conversely, we also observed genes whose transcriptomic expression increased and proteomic expression decreased (**Fig. S3C**). These include *KRT13*, a filament protein known to promote metastasis in BC^51^; *MAFK*, a transcriptional repressor linked to EMT in some breast cancers^52^; *MAP1LC3B2*, a microtubule-associated protein; and *MICAL1*, the actin filament-interacting protein.

To more stringently explore expression trends, we compared significantly downregulated mRNAs to significantly downregulated proteins and found a strong overlap (hypergeometric test, p = 2.05e-163, **Fig. 3D**). The same comparison in significantly upregulated mRNAs and proteins yielded only 3 overlaps: PIP (prolactin-induced protein), CCND1, and HMGCS2 (**Fig. 3E**). PIP has been implicated as essential for proliferation in some luminal A BC cells, and high HMGCS2 levels are associated with tamoxifen resistance^53,54^.

Next, we evaluated whether we observed differential protein expression modulated by alternative splicing changes or cell-cycle dependent transcriptome and proteome changes. Overlap of splicing changes with proteomic changes was minimal but contained key cell cycle factors, including MCM7, INCENP, AURKA and ECT2 (discussed previously) as well as FANCI and FANCA, which coordinate the response to interstrand DNA crosslinks^55^. (**Fig. 3F**). The latter changes suggest loss of the Fanconi Anemia complex function in palbociclib arrested cells, which could influence sensitivity to DNA crosslinking agents. To determine whether the transcripts encoding differentially expressed proteins were controlled in a cell cycle-dependent manner, we compared our dataset to a landmark study characterizing peak gene expression in G1/G1-S and G2 phases^21^. We found overlaps between downregulated proteins in both G1/G1-S and G2-peak mRNA sets, but there remained differentially expressed proteins that were seemingly not captured (**Fig. 3G**, top and bottom left). The same comparison with upregulated proteins showed that only *CCND1* overlapped with the G1/G1-S subset, and only *CCND1* and *TFF3* overlapped with the G2 subset (**Fig. 3G**, top and bottom right). This underscores the consistent and robust Palbo-dependent change to the mRNA- and protein-level expression of cyclin D1. We proceeded to compare our identified up- and down-regulated proteins to 3 external datasets, each identifying proteins with fluctuating cell cycle abundance (**Methods**). Surprisingly, while only 9 proteins were common between all 3 datasets and Palbo-downregulated proteins, all had roles in cell cycle progression (**Fig. S3D**). None of the Palbo-upregulated proteins were common to all datasets, although NDUFAB1 and CCND1 scored in two other datasets^56,57^ (**Fig. S3E**). Overall, this demonstrates the high variability between published proteomics datasets, likely resulting from differences in protein abundance and stability, as these factors vary based on cell type, cellular context and synchronization technique. However, these differences emphasize the importance of studying the effect of cell cycle-targeting therapeutics in clinically relevant cell models that reflect appropriate cancer-types and subtypes. To infer functionality of the differentially regulated proteins, we again employed pathway analysis methods. GO analysis of the downregulated proteins (**Methods**) showed a strong enrichment for cell cycle terms, including mitotic cell cycle process and chromosome organization, further supporting that Palbo suppresses expression of key cell cycle factors (**Fig. 3H**). Consistent with transcriptomic downregulation, we see cell cycle-related terms enriched in the most downregulated proteins (E2F Targets, G2M Checkpoint and Mitotic Spindle) (**Fig. 3I** and **Fig. 3J**, right). Interestingly, we noticed many more enriched terms in the upregulated proteins than in the upregulated genes, suggesting that post-translational changes contribute to these dynamics. While Estrogen Response Early is enriched at both the transcriptomic and proteomic level, we notice several other upregulated terms only at the proteomic level, including Oxidative Phosphorylation, Protein Secretion, and Adipogenesis (**Fig. 3I** and **Fig. 3J**, left). This discordance in the number of upregulated terms between transcripts and protein levels may reflect protein stability; a protein with a long half-life will exhibit constant levels even after it is no longer being transcribed. Additionally, many of the upregulated terms shown in **Fig. 3I** are related to cellular metabolism, and there is a growing body of evidence that suggests CDK4/6i significantly alters various metabolic pathways, including an increase in mitochondrial metabolism^58^.

### Palbociclib induces expression of estrogen-response factors

ER+ breast cancers have long been known to have *CCND1* amplifications and cyclin D1 overexpression^59–62^. As mentioned previously, we saw a significant increase in *CCND1* transcript and cyclin D1 protein levels following Palbo treatment (**Figs. 1B, 3A, 3B**). Additionally, out of all detected cyclin transcripts, the only significantly upregulated cyclin after Palbo treatment was *CCND1* (**Fig. 4A**). These findings, in addition to the importance of estrogen signaling in driving cyclin D1 expression in ER+ BC, motivated us to investigate whether CDK4/6 inhibition affects the expression of ER-target genes and their resulting protein products. As previously described, pathway analysis identified “Estrogen Response Early” as the most significant hallmark term in our upregulated gene set (p-adjusted = 4.53e-04, **Fig. 1D**). Among genes making up the core enrichment subset of the this “Estrogen Response Early” gene set, *CCND1* was the most strongly upregulated and significantly detected member (**Fig. 4B**).

**Figure 4:**
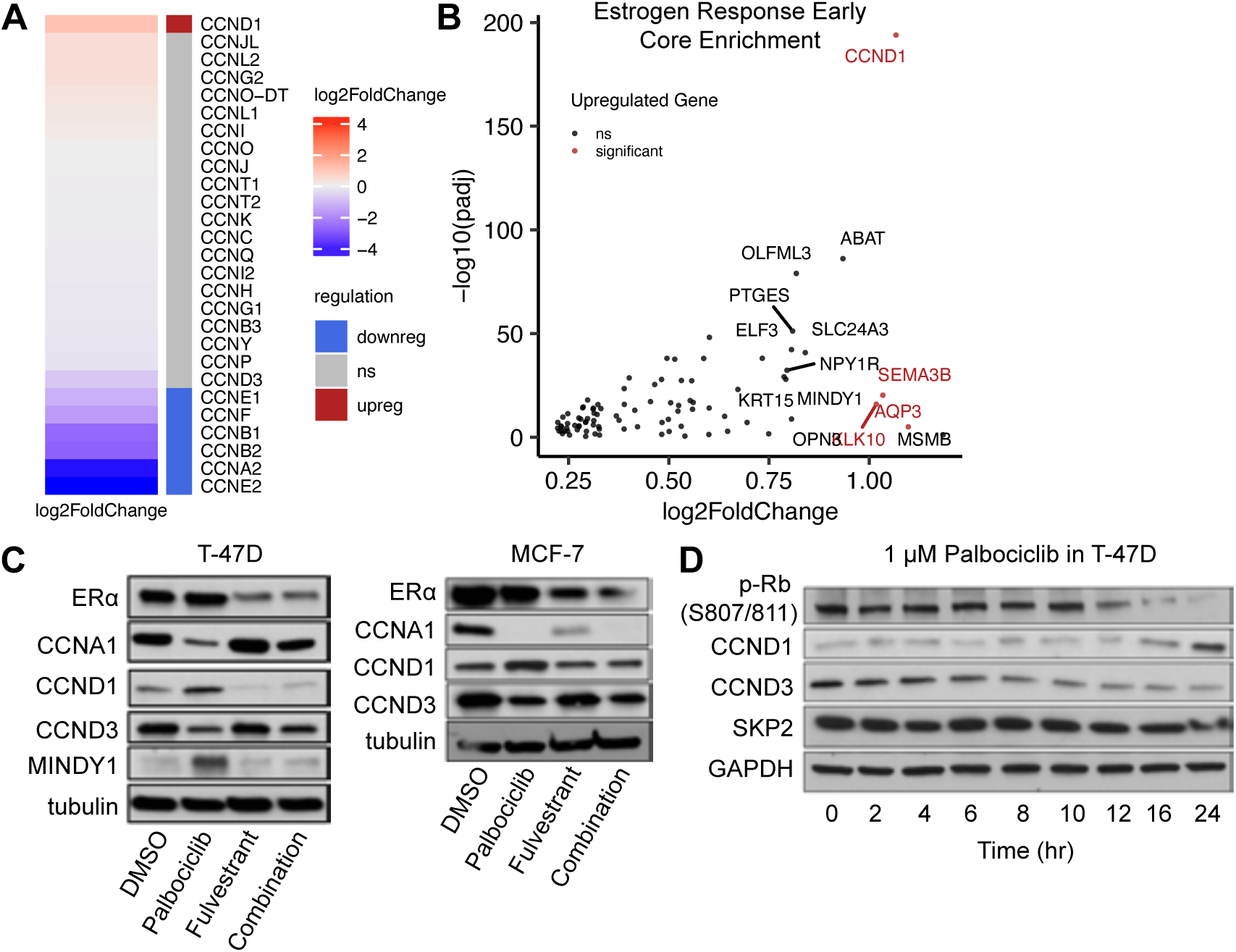
Palbociclib induces expression of estrogen-response factors. **A.** Heat map showing log2FoldChange of all detected cyclin transcripts. Legend: Red: upregulated genes, blue: downregulated genes. **B.** Plot comparing log2FoldChange of genes upregulated by Palbo and genes in the core enrichment subset of the “Estrogen Response Early” gene set from Fig. 1D (top). Red: significantly detected genes passing log2FoldChange ≥ 0.75, black: not significant (ns). **C.** Western blot showing expression of indicated proteins in T-47D (left) and MCF-7 (right) cells after treatment with 1μM Palbo or 1μM fulvestrant alone and in combination for 24 hours. **D.** Western blot showing expression of indicated proteins in T-47D cells treated with 1μM Palbo and collected at the indicated times.

We were surprised to find that Palbo induces an estrogen response because estrogen activates a multitude of factors that ultimately promote cell cycle progression (e.g., *CCND1*). This suggests the presence of under-appreciated negative feedback control between CDK4/6 and ER, wherein CDK4/6 inhibition increases the activity of ER but removes feedback control. Clinically, CDK4/6 inhibitors are given to patients as a combination treatment with anti-hormone therapy, most commonly fulvestrant. This drug works by binding to and inducing the degradation of ER, preventing the activation of its pro-proliferation targets. Therefore, we examined whether the Palbo-dependent upregulation of estrogen response factors persisted in the presence of fulvestrant. We treated T-47D and MCF-7 cells with Palbo or fulvestrant alone and in combination and probed for the indicated proteins by Western blot (**Fig. 4C**). In the presence of Palbo alone, we observed an increase in CCND1 protein, and this effect is abrogated in co-treatment with fulvestrant. In fact, in T-47D cells, fulvestrant resulted in nearly undetectable levels of CCND1. CDK4/6 inhibition with Palbo also increased the expression of MINDY1, another estrogen-dependent gene, and this increase was reversed by co-treatment with fulvestrant. This is particularly interesting as MINDY1 is a deubiquitinase reported to interact with and increase the stability of ERα^63^.

The D-type cyclins (cyclin D1, D2 and D3) are key regulators of the cell cycle, binding to and activating the major G1 kinases, CDK4/6. While initially thought to be functionally redundant, growing evidence indicates that the D-type cyclins can have different expression patterns and distinct functions in different cellular contexts^64,65^. Cyclin D2 is not expressed in most BC cell lines or primary patient tumors^66^; accordingly, in our analysis, we did not detect *CCND2* or cyclin D2 expression. Therefore, we wanted to investigate the expression levels of cyclin D1 and D3. We found that in contrast to cyclin D1, cyclin D3 expression decreases following Palbo treatment in T-47D cells, and we observed this result in another HR+/HER2-BC cell line, MCF-7 (**Fig. 4C**). To further understand the temporal response of cyclin D1 and cyclin D3 expression, we probed for both proteins by Western blot at different times following Palbo exposure in T-47D cells (**Fig. 4D**). Cyclin D1 levels increased at 16 hours and showed significant accumulation at 24 hours, while cyclin D3 levels gradually decreased after Palbo addition. This suggests separate and distinct regulation of the D-type cyclins in response to CDK4/6 inhibition and possibly different functional roles. Collectively, these findings underscore a complex proteomic response to CDK4/6 inhibition within the ER and CDK-RB pathway, which may influence therapeutic responses downstream.

### Palbociclib-responsive genes are enriched for essential genes

Having explored the transcriptomic and proteomic response to CDK 4/6i, we turned to genome-wide CRISPR screening data reported by DepMap ^67^ to better understand the functional consequences of CDK 4/6i-responsive gene regulatory programs. We analyzed gene fitness scores from DepMap across 52 BC cell lines for genes which were differentially expressed following Palbo treatment (**Fig. 5A**, left). These cell lines represent various BC molecular subtypes. Across all cell lines, we extracted and categorized gene fitness/essentiality scores, where a score of –1 or lower indicates greater essentiality and a score close to 0 or higher indicates no essentiality. In total, we extracted 745 differentially expressed genes with available fitness/essentiality scores (**Fig. 5A**, right). We observed that cell lines with similar molecular subtypes, especially among established HR+/HER2-cell lines (T-47D, MCF-7, HCC1428, and ZR751), clustered together based on the fitness scores of Palbo-regulated genes. These data indicate that the responsiveness of ER+/HER2-cells to CDK4/6i is likely dependent on two factors. First is the ability to downregulate a specific set of genes, which would likely be unaffected in other cell lines that, for example, have already dysregulated the CDK/RB pathway. Second is the sensitivity of those cells to the loss of these specific genes/proteins.

**Figure 5:**
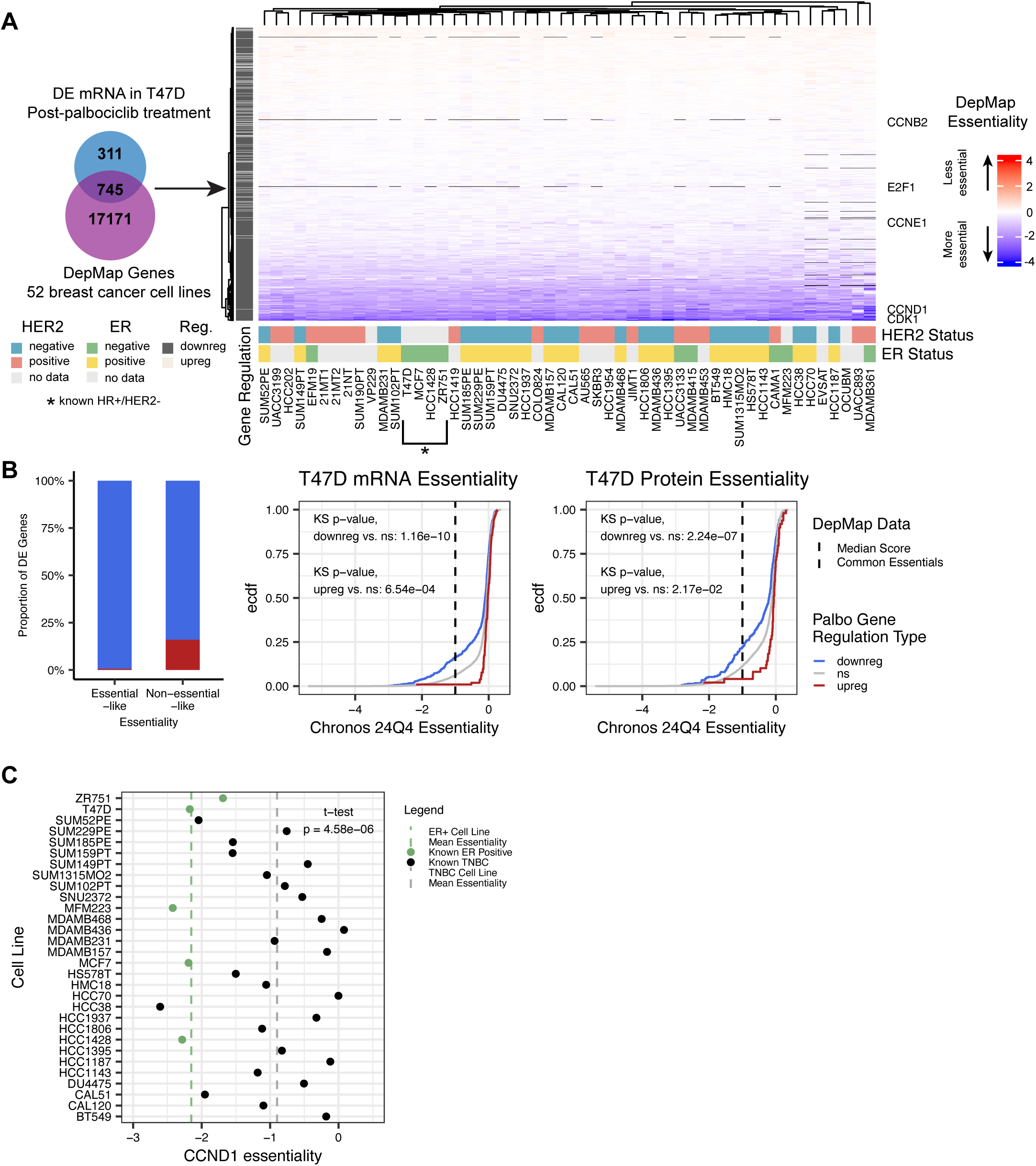
Palbociclib-response genes are enriched for essential genes. **A.** Left: Gene overlap between differentially expressed genes at the mRNA level from Palbo-treated T-47Ds and genes profiled in DepMap across 52 breast cancer cell lines. Right: Heatmap showing gene essentialities of these 745 genes. NA values are represented by black in the heatmap. Bottom annotation: HER2 and ER status obtained from DepMap annotations for each cell line. Left annotation: gene regulation of mRNA post-Palbo treatment. Right annotation: row labels for genes of interest. **B.** Left: proportion of “essential-like” genes and “non-essential-like” DE genes across all 52 cell lines. “Essential-like” genes had essentiality scores ≤ -1. “Non-essential-like” genes had essentiality scores > -1. Middle and right: Cumulative distribution function (CDF) plots of essentiality scores for genes downregulated, upregulated, or not significant (ns) genes after Palbo treatment at the mRNA (middle panel) and protein (right panel) levels. The median score for DepMap common essential genes, -1, is shown by a dashed line. Middle panel: p = 1.16e-10 (downreg vs. ns), p = 6.54e-04 (upreg vs. ns). Right panel: p = 2.24e-07 (downreg vs. ns), p = 2.17e-02 (upreg vs. ns). p-values calculated by Kolmogorov-Smirnov (KS) test. Red line: upregulated genes, blue line: downregulated genes. **C.** CCND1 essentiality scores across for ER+ (green) and TNBC (grey) cell lines. Median essentiality for each group shown by dashed lines. Comparison of mean essentialities: p = 4.58e-06, Student’s t-test.

Building on these observed trends, we determined whether genes that were differentially regulated upon Palbo addition were overall more or less essential than genes that were unaffected. First, we compared the proportions of differentially expressed genes that were more essential or less essential in the T-47D cell line using a fitness score cutoff of –1, the median score of all essential genes (**Fig. 5B**, left). A larger proportion of downregulated genes were found to be essential than upregulated genes. Next, we compared the essentiality score distributions of upregulated, downregulated, and non-significant genes after Palbo treatment and found that downregulated genes are significantly more essential than non-essential genes, both at the mRNA (**Fig. 5B**, middle, p = 1.16e-10, KS test) and protein (**Fig. 5B**, right, p = 2.24e-04, KS test) levels. Intriguingly, upregulated genes were significantly less essential than the unchanged genes both at the mRNA (p = 6.54e-04, KS test) and protein (p = 2.17e-02, KS test) levels.

To verify these observations, we applied a bootstrapping technique wherein we sampled the same number of genes as those that were up- or down-regulated from the list of non-significant genes. We matched based on basal expression level to ensure only genes expressed in T-47D were considered (**Methods**). This “bootstrapping” method yielded randomized control gene sets that could be compared against our up- or down-regulated gene sets. This process was repeated 500 times for both the transcriptomic and proteomic gene sets and confirmed that the “true” set of transcriptionally downregulated genes were consistently more significantly essential than control sets (**Fig. S4A**). The same was true of the proteomic downregulated genes (**Fig. S4B**), indicating that strong essentiality scores are specific to the downregulated genes and not due to random variation. Applying the same procedure to upregulated genes yielded no significant p-adjusted values at the transcriptomic (**Fig. S4C**) or proteomic (**Fig. S4D**) levels. Therefore, while the upregulated genes we observed had weaker essentiality scores than non-essential genes, this does not mean that the upregulated genes are necessarily less essential than the rest of the genes detected as the result was not robust to randomization. Finally, given the prominence of CCND1 upregulation in the Palbo response, we compared CCND1 essentialities across all cell lines surveyed to investigate the relationship between CCND1 and ER+ breast cancers. Comparison of CCND1 essentiality between known ER+ cell lines and known TNBC cell lines from DepMap revealed a significant difference between the two groups (**Fig. 5C**). This is consistent with the cellular response of ER+, but not TNBC, cells to CDK4/6i and the clinical use of CDK4/6i in ER+ BC, but not TNBC ^68^.

### Co-inhibition of CDK4/6 and CDK7 effectively reduces proliferation

Overcoming innate and acquired resistance to targeted therapies, including CDK4/6i, can be informed by understanding the dynamic response to these drugs. Identifying therapeutic combinations can circumvent resistance while also potentially expanding the therapeutic use of CDK4/6 inhibitors to other cancers, where prior clinical trials have been unsuccessful. In clinical treatment of ER+/HER2-BC, palbociclib is administered in conjunction with hormone therapy, and clinical trials testing single agent efficacy in breast cancer have been unsuccessful.

Our comprehensive interrogation of the response to palbociclib in ER+/HER2-BC motivated efforts to test new therapeutic combinations. Notably, our transcriptomic and proteomic data uncovered a negative feedback loop between CDK4/6 and estrogen signaling. We showed that “Estrogen Response Early” is activated by Palbo, and validated that CCND1 and MINDY1 proteins, encoded by ER-regulated genes, are upregulated in response to Palbo. Moreover, their increase was suppressed by fulvestrant. Furthermore, the response to Palbo treatment was widespread. Both G1/S and G2/M genes were downregulated, implying suppression of both RB and DREAM-dependent programs. Thus, re-entry of cells into a proliferative state would need to overcome a deep suppression in the abundance of cell cycle transcripts and proteins, likely requiring activation of one or more CDKs.

A mechanism of CDK4/6-dependent cell cycle control is phosphorylation of the CDK4/6 T-loop by CDK7^69^. CDK7, along with its binding partners MAT1 and cyclin H, comprises the CDK activating kinase complex (CAK). CAK phosphorylates the T-loops of both CDK2 and CDK4/6 to enhance their kinase activity and promote cell cycle progression ^70^. CDK7 also phosphorylates and regulates RNA polymerase II. Interestingly, CDK7, Cyclin H and MAT1 have been reported to increase in ER+ breast cancers at the mRNA and protein level^71^. Further, CDK7 is implicated in the phosphorylation of S118 in ERα, impacting its activation^72^. Additionally, a recent study suggested that CDK7 inhibition could overcome resistance to hormone therapy in ER+/Her2 BC cells^73^.

We therefore reasoned that CDK7 inhibition could be combined with CDK4/6 inhibitors to prevent re-expression of cell cycle proteins and block feedback-dependent activation of ERα. Indeed, a phase one clinical trial shows that the CDK7 inhibitor samuraciclib, in combination with fulvestrant, has anti-tumor activity in HR+/HER2-BC patients whose disease had progressed on CDK4/6i^74^. We determined whether combined inhibition of CDK7 and CDK4/6 (without previous treatment) could additively suppress proliferation. We treated MCF-7 cells with samuraciclib alone or in combination with Palbo. After five days, we performed live cell imaging to obtain a total cell count. We found that Palbo and samuraciclib together decreased cell growth better than either agent alone, with the most significant effect seen at 100nM Palbo addition (**Fig. 6A**). Their combined inhibition suppressed phosphorylation of the CDK2 T-loop (T160) greater than inhibiting either alone. Collectively, this suggests that combining CDK7 and CDK4/6 inhibition could potentially be used to achieve improved therapeutic benefit.

**Figure 6:**
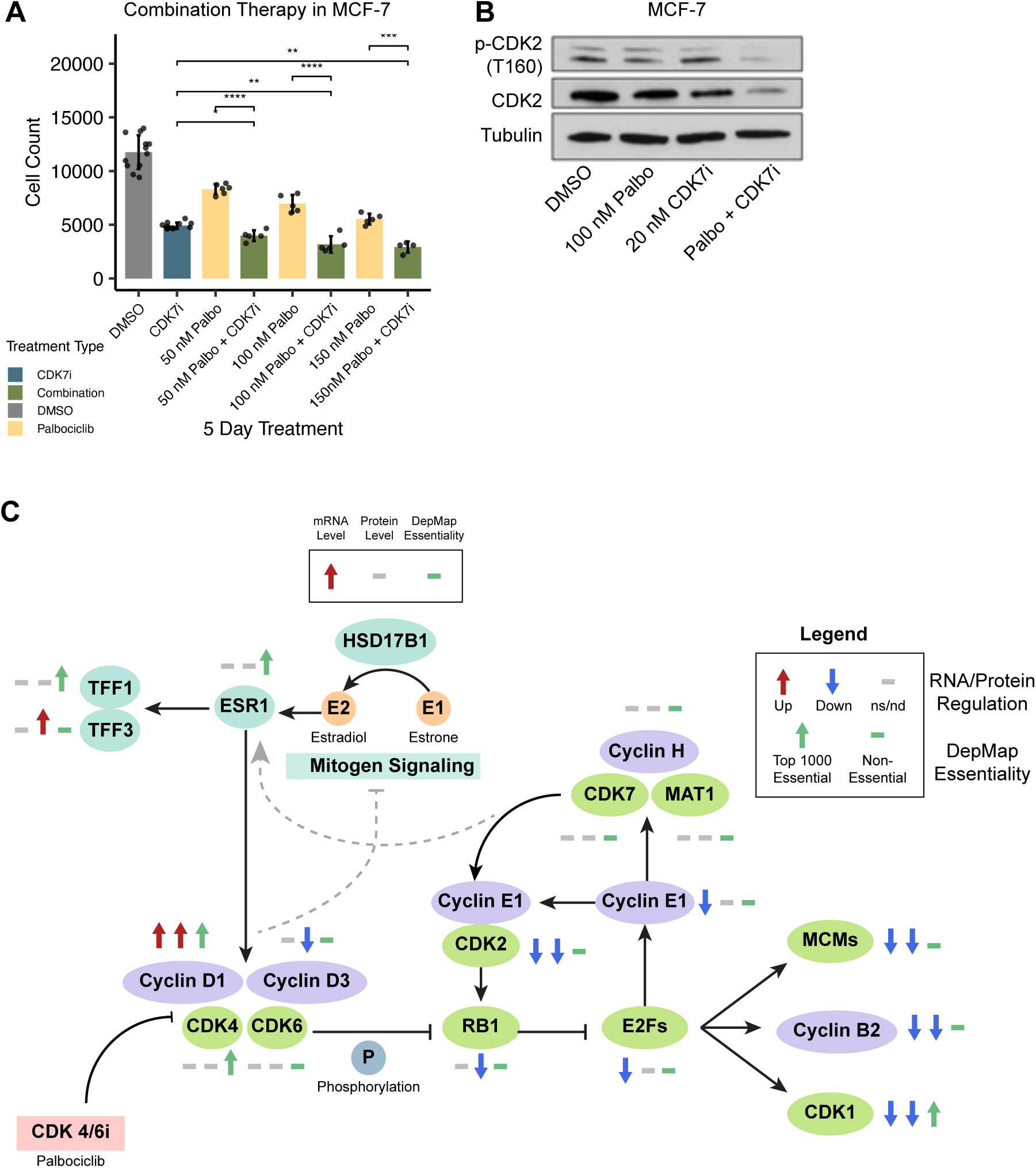
Combined inhibition of CDK4/6 and CDK7. **A:** MCF-7 cell count after 5-day treatment of samuraciclib (CDK7i) or Palbo alone and in combination. **B:** Schematic summarizing pathways described throughout this study, integrating gene and protein regulation and essentiality. The symbols next to each gene/gene family represent observations from the Palbo dataset. Symbol positions: the first position corresponds to transcriptomic differential expression, the second position corresponds to proteomic differential expression, and the third position corresponds to gene essentiality. Symbol colors: Red arrow corresponds to upregulation, blue arrow corresponds to downregulation, gray dash corresponds to no significance (n.s.) or no data available (n.d.). A green arrow at the third represents a gene within the top 1000 T-47D essential genes (z-scored) from DepMap 24Q4 data; a green dash at the third position represents a gene outside of the top 1000 essential T-47D genes.

## DISCUSSION

CDK4/6i have shown significant clinical benefit as a standard-of-care treatment for patients with HR+/HER2-metastatic BC. Still, nearly three decades since the discovery of the Cyclin D-CDK4/6-RB pathway and more than two decades after the development of the first CDK4/6i, patient drug response and resistance represent significant clinical challenges. Surprisingly, we lack a comprehensive molecular understanding of global transcriptional and proteomic changes induced by these inhibitors in BC. How they remodel gene expression, splicing, and proteomics programs in BC remains incompletely understood. To better understand how CDK4/6i affect gene, protein and alternative splicing in cells, we conducted a multi-omics analysis in T-47D BC cells treated with Palbo and observed both broad-patterns and specific gene and protein level changes.

Overall, we found a strong correlation between mRNA and protein changes, and we report a strong Palbo-induced downregulation of genes and their protein products, including *CDK1* and *CDK2, CCNB*2 and *MCM7,* supporting the established mechanism of CDK4/6i to induce a cell cycle arrest via suppression of E2F target genes. In our experiments, we found that many downregulated proteins are also known targets of the proteasomal degradation machinery in cells. This finding supports recent studies that suggest palbociclib can indirectly activate the proteasomal pathway in BC cells to promote a global degradation of proteins while also arresting cells at a time when the Anaphase Promoting Complex/Cyclosome ubiquitin ligase is active towards many cell cycle proteins^75^. We identified a relatively lower number of genes and proteins whose abundance increased after Palbo treatment, and we observed only three upregulated transcripts whose protein product was also upregulated: *PIP*, *CCND1* and *HMGCS2.* From our chromatin accessibility analysis, we report only modest changes induced by Palbo at the 24-hour timepoint. This is consistent with existing studies of CDK4/6i and chromatin remodeling activity^76^.

A novel aspect of this work is our extensive profiling of splicing changes in HR+ BC induced by Palbo treatment. Previous work has demonstrated that mRNA splicing is dynamic throughout the cell cycle^17,21^, and a natural consequence would be that splicing is impacted when the cell cycle is altered by CDK 4/6i. Previous work in melanoma suggests that Palbo inhibits PRMT5, an enzyme that methylates arginine residues, and modulates MDM4 splicing^77^. PRMT5 has emerged as a cancer target in CDK4/6i-resistant ER+ BC^78^, with novel roles in cell cycle progression and post-translational modifications on RNA-binding proteins^79^. In Palbo-resistant HR+ BC models, it has been observed that NSRP1, a splicing regulator, is downregulated^80^. These existing studies underscore the little-explored relationship between RNA processing, cell cycle, and cell cycle inhibition. To our knowledge, we have conducted the first in-depth study of alternative splicing in response to Palbo treatment. In addition to profiling and validating alternative splicing events in candidate cell cycle and RNA processing genes, we identify NMD as a potential post-transcriptional regulatory mechanism shaping the Palbo response. While there has been previous work linking the activity of CDKs to NMD, understanding the relationship of NMD to cell cycle progression, and cell cycle targeted therapies used clinically, has largely been unexplored. NMD effects can be cell-type specific and context-dependent^81^, and thus, to more robustly explore this relationship, further experiments are necessary.

A particularly interesting finding from our studies was the Palbo-induced increase in estrogen signaling (**Fig. 1D**, **Fig. 3J**, **Fig. 6C**). We detected an increase in *HSD17B1* transcripts following Palbo treatment. HSD17B1 plays a key role in estrogen biosynthesis, converting estrone into estradiol E2, a potent ER agonist^82^. This enzyme is also known to promote estrogen signaling^83^. TFF3, an ER target, showed increased levels in response to Palbo. Upregulation of TFF3 promotes cellular proliferation, migration and invasion, and studies have shown that high TFF3 expression correlates with anti-estrogen resistance, providing a link between TFF3 expression and patient clinical outcomes ^84^. In addition, MINDY1 is increased in response to palbociclib, and this has been reported to positively regulate the expression of ERα via deubiquitination. This is consistent with the increase in estrogen response genes following CDH4/6 inactivation, could be part of the feedback loop enforcing ER activity. Altogether, these data provide evidence for a previously undescribed negative feedback loop between CDK4/6 and ER signaling, and they support the notion that combined inhibition of CDK4/6 and other ER-targets may be a suitable avenue to prevent clinical challenges arising from CDK4/6i, including resistance. Our data could also suggest collateral vulnerabilities which arise from either widespread downregulation of the cell cycle proteome or the activation of ER in response to CDK4/6i, with CDK7 representing one attractive example. Finally, our data help clarify why ER+ BC, but not other cancers, are most responsive to CDK4/6i.

## ACKNOWLEDGEMENTS

We thank lab members and colleagues at UNC for helpful discussions throughout this project. The Emanuele lab is supported by the UNC University Cancer Research Fund (UCRF), the National Institutes of Health (R35GM153250 and R01CA280482), and the America Cancer Society (Research Scholar Grant; RSG-18-220-01-TBG). AA was also supported by F31CA288070. The Dominguez lab is supported by NIH R35GM142864 and R01CA290597. AK was supported by T32CA244125. MES was supported by T32GM135095. JCM was supported by NIH 2K12CA120780-16 and the University of North Carolina at Chapel Hill School of Medicine Physician Scientist Training Program. This research is based in part upon work conducted using the UNC Proteomics Core Facility, which is supported in part by NCI Center Core Support Grant (2P30CA016086-45) to the UNC Lineberger Comprehensive Cancer Center. We thank Angie Mordant, Laura Herring, Thomas Webb, Scott Lyons and Natalie Barker for their contributions to the proteomics work performed in this manuscript.

## DATA AVAILABILITY

Sequencing data is available at the Gene Expression Omnibus under series: GSE297939.

## DECLARATION OF INTERESTS

D.D. reports consulting fees from Eve Bio.

## MATERIALS AND METHODS

### Mammalian cell culture, cell lysis, antibodies and reagents

T-47D and MCF-7 cells were obtained from ATCC and grown in DMEM complete medium (Gibco, Cat# 11-965-092) supplemented with 10% fetal bovine serum (EMD Millipore, Cat# F2442). Samples for protein analysis by immunoblot were lysed in NETN buffer [20 mM Tris-HCl (pH 8.0), 100 mM NaCl, 0.5 mM EDTA and 0.5 % (v/v) Nonidet P-40 (NP-40)] supplemented with 2 µg/ml pepstatin, 2 µg/ml apoprotinin, 10 µg/ml leupeptin, 1 mM AEBSF [4-(2 Aminoethyl) benzenesulfonyl fluoride] and 1 mM Na_3_VO_4_. Lysis was performed on ice for 30 minutes with occasional vortexing, and then lysates were spun down at 14,000 rpm for 10 min before determining protein concentration using BCA reagent (Thermo Scientific, Cat# 23227). Standard immunoblotting procedures were followed. A list of reagents and antibodies that were used in this study and the concentrations at which they were used are available in Supplementary Table 5. All antibodies were diluted in 5% nonfat dried milk [diluted in Tris buffered saline, 0.05% tween-20 (TBST)], incubated overnight at 4°C and detected using HRP conjugated secondary antibodies (Jackson Immuno Research Laboratories Inc; 1:5000 dilution).

### Total cell count

MCF-7 cells were seeded at a density of 1K cells per well in 96-well plates (REV-96-Well Greiner 655090 Plate) and treated with samuraciclib (CT7001) (n=12) or palbociclib (n=6 each) alone and in combination for 5 days. Cell nuclei were stained with Hoechst dye (1:1000) in PBS and then imaged with Revvity Celigo Image Cytometer [Nexcelom Bioscience Celigo Pro 5 Channel]. The Celigo expression analysis software [Windows 10 Software Version: 5.5.2.0; Application name: Target 1 + 2] was used to obtain a total cell count in the blue, fluorescent channel [Intensity threshold: 255]. P-values were calculated in R using t.test.

### Global mass spectrometry

#### Sample preparation

T-47D cells were treated with DMSO or 1 µM palbociclib for 24 hours, harvested and lysed in urea buffer [8 M urea, 75 mM NaCl, 50 nM Tris-HCl (pH 8.0) and 1 mM EDTA] supplemented with 2 µg/ml pepstatin, 2 µg/ml apoprotinin, 10 µg/ml leupeptin, 1 mM AEBSF and 1 mM Na_3_VO_4_. Samples were snap frozen in liquid nitrogen and spun down at 14,000 rpm for 20 min before determining protein concentration using BSA reagent. Protein lysates (n = 3) were reduced with 5mM dithiothreitol (DTT; Pierce) at 37°C for 45 min and alkylated with 15mM iodoacetamide (IAA; Pierce) for 45 min at room temperature. Samples were then diluted to 1M urea and subjected to digestion with LysC (Wako) at 37°C for 2h and trypsin (Promega) overnight at 37°C at a 1:50 enzyme:protein ratio. The resulting peptides were acidified to 0.5% trifluoroacetic acid (TFA; Pierce) and desalted using Thermo desalting spin columns. Eluates were dried via vacuum centrifugation and peptide concentration was determined via Pierce Quantitative Fluorometric Assay. All samples were normalized to 0.15 µg/µl and subjected to LC-MS/MS analysis.

#### LC-MS/MS Analysis

The samples (2µl) were analyzed by LC-MS/MS using an Ultimate3000 coupled to an Exploris480 mass Spectrometer (Thermo Scientific). A pooled sample was analyzed before and after the sample set. Samples were injected onto an Ion Opticks Aurora C18 (75 μm id × 15 cm, 1.6 μm particle size) and separated over a 120 min method. The gradient for separation consisted of 5–42% mobile phase B at a 250 nl/min flow rate, where mobile phase A was 0.1% formic acid in water and mobile phase B consisted of 0.1% formic acid in 80% ACN. The Exploris480 was operated in product ion scan mode for Data Independent Acquisition (DIA). A full MS scan (m/z 350-1650) was collected; resolution was set to 120,000 with a maximum injection time of 20 ms and automatic gain control (AGC) target of 300%. Following the full MS scan, a product ion scan was collected (30,000 resolution) and consisted of stepped higher collision dissociation (HCD) set to 25.5, 27, 30; AGC target set to 3000%; maximum injection time set to 55 ms; variable precursor isolation windows from 350-1650 m/z.

#### Proteomics Data Processing and Analysis

Raw data files were processed using Spectronaut (v17; Biognosys) and searched against the Uniprot reviewed human database (UP000005640, containing 20,404 entries, downloaded January 2023) ^85^ and the MaxQuant common contaminants database (246 entries) ^86,87^. The following settings were used: enzyme specificity set to trypsin, up to two missed cleavages allowed, cysteine carbamidomethylation set as a fixed modification, methionine oxidation and N-terminal acetylation set as variable modifications. Precision iRT calibration was enabled. A false discovery rate (FDR) of 1% was used to filter all data. Imputation was disabled, and single hit proteins were excluded. Un-paired student’s t-tests were conducted and p-values, FDR-corrected p-values (q-values), along with log2 fold change ratios were calculated in Spectronaut.

The mass spectrometry proteomics data have been deposited to the ProteomeXchange Consortium via the PRIDE partner repository with the dataset identifier PXD064572.

Data are available via ProteomeXchange with identifier PXD064572. Submission details:

**Project Name:** Proteomic analysis of CDK4/6 inhibition in T47D cells

**Project accession:** PXD064572

**Project DOI:** Not applicable

**Reviewer access details:** Log in to the PRIDE website using the following details:

**Project accession:** PXD064572

**Token:** M7vpnLIrffV1

Alternatively, reviewesr can access the dataset by logging in to the PRIDE website using the following account details:

**Password:** UBRbPEsZ1kcu

### RNA Sequencing Library preparation and Data Processing

RNAs were extracted from cells with RNeasy Kit (Qiagen, 74104). RNA libraries were then generated with KAPA mRNA HyperPrep Kit (Roche, KK8580). Libraries were sequenced on an Illumina platform.

The reference transcriptome was built using the GRCh38 primary assembly FASTA and the GRCh38 primary assembly annotation GTF, both obtained from GENCODE v47^88^. A STAR index was generated using these inputs and a splice junction database overhang (sjdbOverhang) of 99 was used to allow maximum overhang to detect splice junctions from 100 nt reads. Paired-end FASTQs of length 100 nt were obtained from stranded RNA-seq. Reads were trimmed using fastp v0.23.2^89^ with adapter detection enabled. Trimmed FASTQs were aligned with STAR v2.7.10b^90^ using the reference transcriptome and STAR index previously described. Option --twopassMode Basic was used to maximize detection of reads to novel splice junctions and end-to-end read alignment was forced. Alignment files (BAMs) were sorted and indexed using samtools v1.16^91^. multiqc v1.25.1^92^ was used to examine data quality.

### Differential Gene Expression Analysis

Gene expression was quantified from trimmed FASTQs using the quant command of salmon v1.10.2^93^. The -- seqBias and -- gcBias flags were passed for Salmon to correct sequence and GC content bias, respectively, and library type was automatically inferred. Differential gene expression analysis was performed using R v.4.4.3 and v4.5.0^94^ and DESeq2 v1.44.0^95^. For pre-filtering, the minimum group size was set to 3 and the minimum counts were set to 10 as described in the DESeq2 vignette. To determine differential expression (DE), upregulated genes were required to meet a cutoff of log2FoldChange ≥1 and p-adjusted ≤ 0.05; downregulated genes were required to meet a cutoff of log2FoldChange ≤ -1 and p-adjusted ≤ 0.05; all other genes were considered non-significant (ns). ENSEMBL gene IDs were mapped to HGNC gene symbols using org.Hs.eg.db 3.20.0.

### Alternative Splicing Analysis

Alternative splicing (AS) events were calculated from BAM files using rMATS-turbo v4.3.0^22^. The GRCh38 primary assembly GTF (described previously) was used as reference. palbociclib-treated samples were compared to their DMSO-treated counterparts. The data was specified as stranded (--libType fr-firststrand), paired, and read length 100nt. Individual counts were output for all AS events. Events were determined from JCEC (junction and exon read counts) outputs. AS event tables for alternative 3’ splice sites (A3SS), alternative 5’ splice sites (A5SS), retained introns (RI) and skipped exons (SE) were concatenated using R v4.4.0 and subset for only those events on chr1-22 and chrX. Detected events were considered significant if meeting the following criteria: at least 60 reads mapping to either inclusion junction reads or skipped junction reads (10 reads per sample), a false discovery rate (FDR) ≤ 0.05, and abs(Inclusion Level Difference) ≥ 0.05. High-confidence SE and RI events were selected for splicing validation if meeting the following criteria: at least 300 reads mapping to either inclusion junction reads or skipped junction reads, an FDR ≤ 0.05, abs(Inclusion Level Difference) ≥ 0.1, and inclusion form length ≤ 1000 to limit the size of the product for validation.

### Proteomics Analysis

Proteomics data was processed in R v4.4.3. Genes were classified as upregulated if meeting an average log2FoldChange (AVG.Log2.Ratio) ≥ 0.6 and a Qvalue ≤ 0.05, downregulated if meeting an average log2FoldChange ≤ -0.6 and a Qvalue ≤ 0.05, and non-significant (ns) otherwise. Non-human genes were removed and ENSEMBL gene IDs were mapped from peptide IDs (UniProt) or HGNC gene symbols (if available) using org.Hs.eg.db 3.20.0 and collated. Proteins with no available gene ID mapping were excluded from analysis. The mapped gene IDs corresponding to proteins were used for all downstream analyses.

### ATAC Procedure

We adapted the ATAC-seq protocol from the original methods described by Buenrostro et al.^96^ and Corces et al.^97^. We cultured T-47D cells from three distinct biological replicates in separate 60 mm dishes and allowed them to reach 60-70% confluency. We replaced the culture media (DMEM with 10% FBS, Penicillin, and Streptomycin) entirely with media containing either 0.1% DMSO (control) or 1 µM palbociclib and incubated for 24 hours prior to the ATAC assay.

We washed the cells three times with ice-cold PBS, then incubated them with 0.25% Trypsin-EDTA for 5 minutes. We quenched the trypsin with complete media (twice the volume) and centrifuged the cells at 500 × g for 5 minutes. Cells were washed twice more with ice-cold PBS, resuspended in ATAC resuspension buffer (RSP; 10 mM Tris-HCl, 10 mM NaCl, 3 mM MgCl2), and counted.

We aliquoted 50,000 cells per biological replicate, centrifuged them, and resuspended the pellets in lysis buffer (RSP supplemented with 0.1% Tween-20, 0.1% NP-40, and 0.01% Digitonin) for a 5-minute incubation. We added 1 mL of wash buffer (RSP with 0.1% Tween-20), centrifuged the cells again, and resuspended them in tagmentation buffer (Diagenode 2× buffer [C01019043] diluted to 1×, supplemented with 0.01% Digitonin, 0.1% Tween-20, and 2.5 µL Diagenode Tn5 enzyme [C01070010]). We incubated the reaction for 30 minutes at 37°C. We purified the DNA using the Zymo DNA Clean & Concentrator kit, following the manufacturer’s instructions, and eluted in 10 µL. For library amplification, we used the NEBNext® High-Fidelity 2X PCR Master Mix (M0541) and Diagenode UDI Tagmentation Library Set (C01011034) following this thermal cycling protocol: 72°C for 5 minutes, 98°C for 30 seconds, 10 cycles of: 98°C for 10 seconds, 63°C for 30 seconds, and one last cycle of 72°C for 1 minute. We size-selected libraries within a range of 200-700 bp using SPRISelect (B23317) beads, pooled the libraries, and sequenced them using a NextSeq 1000 platform.

### ATAC-seq Data Processing and Analysis

We first assessed the quality of paired-end ATAC-seq reads using FastQC (v0.12.1-0)^98^. To reduce alignment noise from non-nuclear sources, we employed a serial pre-alignment strategy. We initially aligned reads to a ribosomal DNA (rtDNA) index using Bowtie2 (v2.5.4-6)^99^ with the parameters --local --very-sensitive-local. Reads that did not align to rtDNA were subsequently aligned to the mitochondrial genome (chrM) using the same parameters. The remaining unaligned reads, representing the clean nuclear fraction, proceeded to genomic alignment.

We aligned the clean paired-end reads to the GRCh38 human reference genome assembly using Bowtie2 with parameters --local --very-sensitive-local-X 2000. We piped these alignments to Samtools (v1.22.1-0)^100^ to filter for properly paired reads with a mapping quality score (MAPQ) of at least 30, while excluding supplementary alignments, unmapped reads, and secondary alignments (flag 1804). We then position-sorted the filtered alignments and identified and removed PCR duplicates using Picard (v3.4.0-0) MarkDuplicates^101^ with the REMOVE_DUPLICATES=true option. We indexed the final deduplicated BAM files with Samtools. Throughout the pipeline, we collected quality control metrics, including alignment statistics with Samtools flagstat and stats, and fragment insert size distributions with Picard CollectInsertSizeMetrics.

For downstream analysis, we adjusted alignment start sites to center reads on the Tn5 transposase cutting site. We performed this step using the alignmentSieve command from deepTools (v3.5.6-0)^102^ with the -- ATACshift option. We then sorted the resulting Tn5-shifted BAM files by position and indexed them with Samtools.

We performed peak calling using two methods within the MACS3 (v3.0.3-0)^103^ framework. First, we called peaks on the Tn5-shifted BAM files using the macs3 callpeak command with a sq-value cutoff of 0.05. We specified the --nomodel option along with --shift -75 and --extsize 150 to define the nucleosome-free region model based on the known Tn5 insertion offset. In parallel, we ran the macs3 hmmratac command on the original (unshifted) deduplicated BAM files to utilize its Hidden Markov Model for identifying accessible chromatin regions. For both methods, we filtered the resulting peak sets to exclude regions listed in the ENCODE hg38 blacklist (v2).

Resulting IDR BED files were merged using bedtools v2.31^104^ and converted to SAF format. Counts were calculated using the featureCounts option from subread v2.0.6^105,106^ to count chromatin accessibility peaks. Differential chromatin accessibility was calculated using DESeq2 and filtered according to vignette specifications. Genomic regions for mRNAs were calculated using a TxDb object built from GRCh38 (GENCODE V47). 1 kb upstream was added to all mRNA regions using the GenomicRanges resize function. mRNA regions were overlapped with chromatin accesibility regions using the GenomicRanges findOverlaps function.

### Additional Bioinformatic Analysis and Integration

#### Gene Ontology Analysis

Transcriptomic and proteomic analysis was performed in R v4.4.0 using the enrichGO function from clusterProfiler v4.14.6^107^ using downregulated genes/proteins as the gene set and all detected genes/proteins as the universe to provide a comparable background. Ontology “BP” (biological process) was used, and p- and q-value significance cutoffs were set to 0.05.

#### Gene Set Enrichment Analysis

Transcriptomic and proteomic analysis was performed in R v4.4.0 using the GSEA function from clusterProfiler v4.14.6^108^ (internally by fgsea v1.32.4^109^). Genes/proteins were ranked by score = log2FoldChange * (1 - p-value). In cases of duplicated genes, the gene with the lower absolute score was dropped. Hallmark gene sets were obtained from MSigDb^110^. An abs(NES) of 1.5 and a p-adjusted value of 0.05 were used for gene set significance cutoffs.

#### Periodic Gene Expression (mRNA)

Supplemental data were obtained and processed from the following publications for transcriptomic analysis: Dominguez et al 2016^18^ Table S1, periodic gene expression; Li, Wang & Wang et al 2022 ^20^ Table S4, periodic transcripts across 3 cancer cell lines (only transcripts mappable to ENSEMBL gene IDs were used for analysis); Boström et al 2017^19^ Table S2, periodic gene expression (mapped to ENSEMBL gene IDs for analysis); Whitfield et al 2002^21^.

#### Periodic Splicing

Raw FASTQs from Dominguez et al 2016^17^ were reprocessed. BAMs were generated using STAR via alignment to GRCh38 (GENCODE v45) and a custom GTF file modified from GENCODE 45. Splicing analysis was carried out in rMATS. UniProt was queried using the R library drawProteins v1.26.0^111^.

#### SMG KD Overlap

RNA-seq FASTQs from SMG6 and SMG7 KD MCF-7 cells was obtained from Britto-Borges et al. 2024^41^. Reads were trimmed and preprocessed using fastp v0.23.2^89^ and aligned and sorted using STAR v2.7.11b^90^ and samtools v1.22^91^. Alternative splicing (AS) events were calculated from BAM files using rMATS-turbo v4.3.0^22^. The GRCh38 primary assembly GTF (described previously) was used as reference. As with the palbociclib splicing analysis, events were determined from JCEC (junction and exon read counts) outputs. AS event tables for A3SS, A5SS, SE, and RI events were concatenated using R v4.5.0 and subset for only those events on chr1-22 and chrX. Detected events were considered significant if meeting the following criteria: at least 300 reads mapping to either inclusion junction reads or skipped junction reads (50 reads per sample), a false discovery rate (FDR) ≤ 0.05, and abs(Inclusion Level Difference) ≥ 0.05.

#### Periodic Gene Expression (Protein)

Supplemental data were obtained and processed from the following publications for proteomic analysis: Ly et al 2014^57^ Table S1 (genes were included if detected in asynchronous conditions, present in all fractionations, and labeled as cell-cycle regulated in Table S3); Mahdessian et al 2021 ^56^ Table S2 (interphase genes labeled as CCD proteins); Lane et al 2013^112^ Table S1 (genes mapped to ENSEMBL IDs from UniProt IDs).

#### DepMap Analysis

CRISPR Gene Effect Score tables and Model ID metadata were downloaded from the DepMap 24Q4 release^113^ and subset for breast tissue. Cell lines were excluded from analysis if labeled as “non-cancerous” or as a cell line subclone in the metadata. NA gene essentialities were not excluded.

### Splicing Validation

#### Primer design

Primers for SMARCA1, MCM7, AURKA, INCENP, ECT2, and DHX34 were designed using the UCSC Genome Browser (hg38) In-Silico PCR tool^114^ and primer3^115^. Primers for all splicing events shown are available in Supplementary Table 7.

#### Sample preparation

6 biological replicates of T-47D cells were treated with 1µM palbociclib or equal volume 100% DMSO for 24 hours. In addition, for NMD validation 3 biological replicates of T-47D cells were treated with 1µM SMG1 inhibitor, palbociclib, or abemaciclib for 24 hours or 100µg/ml cycloheximide for 6 hours. Cells were pelleted and collected. RNA was extracted from cell pellets using RNeasy kit with DNase I (Qiagen, 74106).

#### RT-PCR

cDNA was synthesized from RNA extracted above using Maxima Master Mix (Thermo Fisher, M1662). RT-PCR was performed using Q5® Hot Start High-Fidelity 2X Master Mix (New England Biolabs, M0494S). Samples were run on 3% agarose (Invitrogen, 16500-500) TAE (Fisher Scientific, BP13324) gel. INCENP was run on Novex™ TBE Gels, 10% (Invitrogen, EC63752BOX). PSI values were quantified using ImageJ and Student’s two-way t-tests were performed using Graphpad Prism 10. Reagents used for PCR: 1.25uM forward/reverse primer, 170ng cDNA, 1x Q5 Master mix. Program: Initial melt 15s 98°C followed by 28 cycles of melt:30s@98°C, anneal:30s@65°C, extend:30-60s 72°C, and a final extension of 4 min@72°C. After the following reactions, samples were held at 4°C.

### Plotting and statistical analyses

Unless otherwise described, statistical analysis and plotting was done in R v4.4.3 and 4.5.0. Similarly, unless otherwise described, all tests used for statistical analyses are noted in the main text and figures. Attached R packages to carry out analyses are listed in Supplementary Table 5. Figures were assembled in Adobe Illustrator.

## REFERENCES

(1) Sherman, R. L.; Firth, A. U.; Henley, S. J.; Siegel, R. L.; Negoita, S.; Sung, H.; Kohler, B. A.; Anderson, R. N.; Cucinelli, J.; Scott, S.; Benard, V. B.; Richardson, L. C.; Jemal, A.; Cronin, K. A. Annual Report to the Nation on the Status of Cancer, Featuring State-Level Statistics after the Onset of the COVID-19 Pandemic. Cancer 2025, 131 (9), e35833. 10.1002/cncr.35833.

(2) Wu, L.; Timmers, C.; Maiti, B.; Saavedra, H. I.; Sang, L.; Chong, G. T.; Nuckolls, F.; Giangrande, P.; Wright, F. A.; Field, S. J.; Greenberg, M. E.; Orkin, S.; Nevins, J. R.; Robinson, M. L.; Leone, G. The E2F1-3 Transcription Factors Are Essential for Cellular Proliferation. Nature 2001, 414 (6862), 457–462. 10.1038/35106593.

(3) Chellappan, S. P.; Hiebert, S.; Mudryj, M.; Horowitz, J. M.; Nevins, J. R. The E2F Transcription Factor Is a Cellular Target for the RB Protein. Cell 1991, 65 (6), 1053–1061. 10.1016/0092-8674(91)90557-f.

(4) Lees, J. A.; Saito, M.; Vidal, M.; Valentine, M.; Look, T.; Harlow, E.; Dyson, N.; Helin, K. The Retinoblastoma Protein Binds to a Family of E2F Transcription Factors. Mol Cell Biol 1993, 13 (12), 7813–7825. 10.1128/mcb.13.12.7813-7825.1993.

(5) Calzone, L.; Gelay, A.; Zinovyev, A.; Radvanyi, F.; Barillot, E. A Comprehensive Modular Map of Molecular Interactions in RB/E2F Pathway. Mol Syst Biol 2008, 4, 173. 10.1038/msb.2008.7.

(6) Morrison, L.; Loibl, S.; Turner, N. C. The CDK4/6 Inhibitor Revolution - a Game-Changing Era for Breast Cancer Treatment. Nat Rev Clin Oncol 2024, 21 (2), 89–105. 10.1038/s41571-023-00840-4.

(7) Turner, N. C.; Ro, J.; André, F.; Loi, S.; Verma, S.; Iwata, H.; Harbeck, N.; Loibl, S.; Huang Bartlett, C.; Zhang, K.; Giorgetti, C.; Randolph, S.; Koehler, M.; Cristofanilli, M. Palbociclib in Hormone-Receptor–Positive Advanced Breast Cancer. New England Journal of Medicine 2015, 373 (3), 209–219. 10.1056/nejmoa1505270.

(8) Harbeck, N.; Brufsky, A.; Rose, C. G.; Korytowsky, B.; Chen, C.; Tantakoun, K.; Jazexhi, E.; Nguyen, D. H. V.; Bartlett, M.; Samjoo, I. A.; Pluard, T. Real-World Effectiveness of CDK4/6i in First-Line Treatment of HR+/HER2-Advanced/Metastatic Breast Cancer: Updated Systematic Review. Front Oncol 2025, 15, 1530391. 10.3389/fonc.2025.1530391.

(9) Pu, D.; Xu, D.; Wu, Y.; Chen, H.; Shi, G.; Feng, D.; Zhang, M.; Liu, Z.; Li, J. Efficacy of CDK4/6 Inhibitors Combined with Endocrine Therapy in HR+/HER2-Breast Cancer: An Umbrella Review. J Cancer Res Clin Oncol 2024, 150 (1), 16. 10.1007/s00432-023-05516-1.

(10) Rinnerthaler, G.; Gampenrieder, S. P.; Greil, R. ASCO 2018 Highlights: Metastatic Breast Cancer. Memo 2018, 11 (4), 276–279. 10.1007/s12254-018-0450-9.

(11) McCartney, A.; Migliaccio, I.; Bonechi, M.; Biagioni, C.; Romagnoli, D.; De Luca, F.; Galardi, F.; Risi, E.; De Santo, I.; Benelli, M.; Malorni, L.; Di Leo, A. Mechanisms of Resistance to CDK4/6 Inhibitors: Potential Implications and Biomarkers for Clinical Practice. Front Oncol 2019, 9, 666. 10.3389/fonc.2019.00666.

(12) Xu, X.-Q.; Pan, X.-H.; Wang, T.-T.; Wang, J.; Yang, B.; He, Q.-J.; Ding, L. Intrinsic and Acquired Resistance to CDK4/6 Inhibitors and Potential Overcoming Strategies. Acta Pharmacol Sin 2021, 42 (2), 171–178. 10.1038/s41401-020-0416-4.

(13) Hopkins, B. D.; Pauli, C.; Du, X.; Wang, D. G.; Li, X.; Wu, D.; Amadiume, S. C.; Goncalves, M. A.; Hodakoski, C.; Lundquist, M. R.; Bareja, R.; Ma, Y.; Harris, E. M.; Sboner, A.; Beltran, H.; Rubin, M. A.; Mukherjee, S.; Cantley, L. C. Suppression of Insulin Feedback Enhances the Efficacy of PI3K Inhibitors. Nature 2018, 560 (7719), 499–503. 10.1038/s41586-018-0343-4.

(14) Love, M. I.; Huber, W.; Anders, S. Moderated Estimation of Fold Change and Dispersion for RNA-Seq Data with DESeq2. Genome Biol 2014, 15 (12), 550. 10.1186/s13059-014-0550-8.

(15) Subramanian, A.; Tamayo, P.; Mootha, V. K.; Mukherjee, S.; Ebert, B. L.; Gillette, M. A.; Paulovich, A.; Pomeroy, S. L.; Golub, T. R.; Lander, E. S.; Mesirov, J. P. Gene Set Enrichment Analysis: A Knowledge-Based Approach for Interpreting Genome-Wide Expression Profiles. Proc Natl Acad Sci U S A 2005, 102 (43), 15545–15550. 10.1073/pnas.0506580102.

(16) Spellman, P. T.; Sherlock, G.; Zhang, M. Q.; Iyer, V. R.; Anders, K.; Eisen, M. B.; Brown, P. O.; Botstein, D.; Futcher, B. Comprehensive Identification of Cell Cycle-Regulated Genes of the Yeast Saccharomyces Cerevisiae by Microarray Hybridization. Mol Biol Cell 1998, 9 (12), 3273–3297. 10.1091/mbc.9.12.3273.

(17) Dominguez, D.; Tsai, Y.-H.; Weatheritt, R.; Wang, Y.; Blencowe, B. J.; Wang, Z. An Extensive Program of Periodic Alternative Splicing Linked to Cell Cycle Progression. Elife 2016, 5. 10.7554/eLife.10288.

(18) Dominguez, D.; Tsai, Y.-H.; Gomez, N.; Jha, D. K.; Davis, I.; Wang, Z. A High-Resolution Transcriptome Map of Cell Cycle Reveals Novel Connections between Periodic Genes and Cancer. Cell Res 2016, 26 (8), 946–962. 10.1038/cr.2016.84.

(19) Boström, J.; Sramkova, Z.; Salašová, A.; Johard, H.; Mahdessian, D.; Fedr, R.; Marks, C.; Medalová, J.; Souček, K.; Lundberg, E.; Linnarsson, S.; Bryja, V.; Sekyrova, P.; Altun, M.; Andäng, M. Comparative Cell Cycle Transcriptomics Reveals Synchronization of Developmental Transcription Factor Networks in Cancer Cells. PLoS One 2017, 12 (12), e0188772. 10.1371/journal.pone.0188772.

(20) Li, C.-X.; Wang, J.-S.; Wang, W.-N.; Xu, D.-K.; Zhou, Y.-T.; Sun, F.-Z.; Li, Y.-Q.; Guo, F.-Z.; Ma, J.-L.; Zhang, X.-Y.; Chang, M.-J.; Xu, B.-H.; Ma, F.; Qian, H.-L. Expression Dynamics of Periodic Transcripts during Cancer Cell Cycle Progression and Their Correlation with Anticancer Drug Sensitivity. Mil Med Res 2022, 9 (1), 71. 10.1186/s40779-022-00432-w.

(21) Whitfield, M. L.; Sherlock, G.; Saldanha, A. J.; Murray, J. I.; Ball, C. A.; Alexander, K. E.; Matese, J. C.; Perou, C. M.; Hurt, M. M.; Brown, P. O.; Botstein, D. Identification of Genes Periodically Expressed in the Human Cell Cycle and Their Expression in Tumors. Mol Biol Cell 2002, 13 (6), 1977–2000. 10.1091/mbc.02-02-0030.

(22) Shen, S.; Park, J. W.; Lu, Z.; Lin, L.; Henry, M. D.; Wu, Y. N.; Zhou, Q.; Xing, Y. RMATS: Robust and Flexible Detection of Differential Alternative Splicing from Replicate RNA-Seq Data. Proc Natl Acad Sci U S A 2014, 111 (51), E5593–601. 10.1073/pnas.1419161111.

(23) Middleton, R.; Gao, D.; Thomas, A.; Singh, B.; Au, A.; Wong, J. J.-L.; Bomane, A.; Cosson, B.; Eyras, E.; Rasko, J. E. J.; Ritchie, W. IRFinder: Assessing the Impact of Intron Retention on Mammalian Gene Expression. Genome Biol 2017, 18 (1), 51. 10.1186/s13059-017-1184-4.

(24) Rao, G. H.; White, J. G. An Improved Method for Measuring Endogenous Serotonin in Platelets of Patients with Hermansky-Pudlak Syndrome. Thromb Res 1988, 51 (2), 225–227. 10.1016/0049-3848(88)90066-7.

(25) Ni, T.; Yang, W.; Han, M.; Zhang, Y.; Shen, T.; Nie, H.; Zhou, Z.; Dai, Y.; Yang, Y.; Liu, P.; Cui, K.; Zeng, Z.; Tian, Y.; Zhou, B.; Wei, G.; Zhao, K.; Peng, W.; Zhu, J. Global Intron Retention Mediated Gene Regulation during CD4+ T Cell Activation. Nucleic Acids Res 2016, 44 (14), 6817–6829. 10.1093/nar/gkw591.

(26) Pan, Q.; Shai, O.; Misquitta, C.; Zhang, W.; Saltzman, A. L.; Mohammad, N.; Babak, T.; Siu, H.; Hughes, T. R.; Morris, Q. D.; Frey, B. J.; Blencowe, B. J. Revealing Global Regulatory Features of Mammalian Alternative Splicing Using a Quantitative Microarray Platform. Mol Cell 2004, 16 (6), 929–941. 10.1016/j.molcel.2004.12.004.

(27) Damodaran, A. P.; Gavard, O.; Gagné, J.-P.; Rogalska, M. E.; Behera, A. K.; Mancini, E.; Bertolin, G.; Courtheoux, T.; Kumari, B.; Cailloce, J.; Mereau, A.; Poirier, G. G.; Valcárcel, J.; Gonatopoulos-Pournatzis, T.; Watrin, E.; Prigent, C. Proteomic Study Identifies Aurora-A-Mediated Regulation of Alternative Splicing through Multiple Splicing Factors. J Biol Chem 2025, 301 (1), 108000. 10.1016/j.jbc.2024.108000.

(28) Sasai, K.; Katayama, H.; Hawke, D. H.; Sen, S. Aurora-C Interactions with Survivin and INCENP Reveal Shared and Distinct Features Compared with Aurora-B Chromosome Passenger Protein Complex. PLoS One 2016, 11 (6), e0157305. 10.1371/journal.pone.0157305.

(29) Honda, R.; Körner, R.; Nigg, E. A. Exploring the Functional Interactions between Aurora B, INCENP, and Survivin in Mitosis. Mol Biol Cell 2003, 14 (8), 3325–3341. 10.1091/mbc.e02-11-0769.

(30) Samejima, K.; Platani, M.; Wolny, M.; Ogawa, H.; Vargiu, G.; Knight, P. J.; Peckham, M.; Earnshaw, W. C. The Inner Centromere Protein (INCENP) Coil Is a Single α-Helix (SAH) Domain That Binds Directly to Microtubules and Is Important for Chromosome Passenger Complex (CPC) Localization and Function in Mitosis. J Biol Chem 2015, 290 (35), 21460–21472. 10.1074/jbc.M115.645317.

(31) Honda, R.; Körner, R.; Nigg, E. A. Exploring the Functional Interactions between Aurora B, INCENP, and Survivin in Mitosis. Mol Biol Cell 2003, 14 (8), 3325–3341. 10.1091/mbc.e02-11-0769.

(32) Li, D.; Wang, X.; Miao, H.; Liu, H.; Pang, M.; Guo, H.; Ge, M.; Glass, S. E.; Emmrich, S.; Ji, S.; Zhou, Y.; Ye, X.; Mao, H.; Wang, J.; Liu, Q.; Kim, T.; Klusmann, J.-H.; Li, C.; Liu, Z.; Jin, H.; Nie, Y.; Wu, K.; Fan, D.; Song, X.; Wang, X.; Li, L.; Lu, Y.; Zhao, X. The LncRNA MIR99AHG Directs Alternative Splicing of SMARCA1 by PTBP1 to Enable Invadopodia Formation in Colorectal Cancer Cells. Sci Signal 2023, 16 (803), eadh4210. 10.1126/scisignal.adh4210.

(33) Barak, O.; Lazzaro, M. A.; Cooch, N. S.; Picketts, D. J.; Shiekhattar, R. A Tissue-Specific, Naturally Occurring Human SNF2L Variant Inactivates Chromatin Remodeling. J Biol Chem 2004, 279 (43), 45130–45138. 10.1074/jbc.M406212200.

(34) Tatsumoto, T.; Xie, X.; Blumenthal, R.; Okamoto, I.; Miki, T. Human ECT2 Is an Exchange Factor for Rho GTPases, Phosphorylated in G2/M Phases, and Involved in Cytokinesis. J Cell Biol 1999, 147 (5), 921–928. 10.1083/jcb.147.5.921.

(35) Lewis, B. P.; Green, R. E.; Brenner, S. E. Evidence for the Widespread Coupling of Alternative Splicing and Nonsense-Mediated MRNA Decay in Humans. Proc Natl Acad Sci U S A 2003, 100 (1), 189–192. 10.1073/pnas.0136770100.

(36) Evrin, C.; Clarke, P.; Zech, J.; Lurz, R.; Sun, J.; Uhle, S.; Li, H.; Stillman, B.; Speck, C. A Double-Hexameric MCM2-7 Complex Is Loaded onto Origin DNA during Licensing of Eukaryotic DNA Replication. Proc Natl Acad Sci U S A 2009, 106 (48), 20240–20245. 10.1073/pnas.0911500106.

(37) Hug, N.; Cáceres, J. F. The RNA Helicase DHX34 Activates NMD by Promoting a Transition from the Surveillance to the Decay-Inducing Complex. Cell Rep 2014, 8 (6), 1845–1856. 10.1016/j.celrep.2014.08.020.

(38) Wong, J. J.-L.; Ritchie, W.; Ebner, O. A.; Selbach, M.; Wong, J. W. H.; Huang, Y.; Gao, D.; Pinello, N.; Gonzalez, M.; Baidya, K.; Thoeng, A.; Khoo, T.-L.; Bailey, C. G.; Holst, J.; Rasko, J. A. J. Orchestrated Intron Retention Regulates Normal Granulocyte Differentiation. Cell 2013, 154 (3), 583–595. 10.1016/j.cell.2013.06.052.

(39) Yamashita, A.; Ohnishi, T.; Kashima, I.; Taya, Y.; Ohno, S. Human SMG-1, a Novel Phosphatidylinositol 3-Kinase-Related Protein Kinase, Associates with Components of the MRNA Surveillance Complex and Is Involved in the Regulation of Nonsense-Mediated MRNA Decay. Genes Dev 2001, 15 (17), 2215–2228. 10.1101/gad.913001.

(40) Gopalsamy, A.; Bennett, E. M.; Shi, M.; Zhang, W.-G.; Bard, J.; Yu, K. Identification of Pyrimidine Derivatives as HSMG-1 Inhibitors. Bioorg Med Chem Lett 2012, 22 (21), 6636–6641. 10.1016/j.bmcl.2012.08.107.

(41) Britto-Borges, T.; Gehring, N. H.; Boehm, V.; Dieterich, C. NMDtxDB: Data-Driven Identification and Annotation of Human NMD Target Transcripts. RNA 2024, 30 (10), 1277–1291. 10.1261/rna.080066.124.

(42) Britto-Borges, T.; Gehring, N. H.; Boehm, V.; Dieterich, C. NMDtxDB: Data-Driven Identification and Annotation of Human NMD Target Transcripts. RNA 2024, 30 (10), 1277–1291. 10.1261/rna.080066.124.

(43) Rega, C.; Tsitsa, I.; Roumeliotis, T. I.; Krystkowiak, I.; Portillo, M.; Yu, L.; Vorhauser, J.; Pines, J.; Mansfeld, J.; Choudhary, J.; Davey, N. E. High Resolution Profiling of Cell Cycle-Dependent Protein and Phosphorylation Abundance Changes in Non-Transformed Cells. Nat Commun 2025, 16 (1), 2579. 10.1038/s41467-025-57537-8.

(44) Franks, J. L.; Martinez-Chacin, R. C.; Wang, X.; Tiedemann, R. L.; Bonacci, T.; Choudhury, R.; Bolhuis, D. L.; Enrico, T. P.; Mouery, R. D.; Damrauer, J. S.; Yan, F.; Harrison, J. S.; Major, M. Ben; Hoadley, K. A.; Suzuki, A.; Rothbart, S. B.; Brown, N. G.; Emanuele, M. J. In Silico APC/C Substrate Discovery Reveals Cell Cycle-Dependent Degradation of UHRF1 and Other Chromatin Regulators. PLoS Biol 2020, 18 (12), e3000975. 10.1371/journal.pbio.3000975.

(45) Michowski, W.; Chick, J. M.; Chu, C.; Kolodziejczyk, A.; Wang, Y.; Suski, J. M.; Abraham, B.; Anders, L.; Day, D.; Dunkl, L. M.; Li Cheong Man, M.; Zhang, T.; Laphanuwat, P.; Bacon, N. A.; Liu, L.; Fassl, A.; Sharma, S.; Otto, T.; Jecrois, E.; Han, R.; Sweeney, K. E.; Marro, S.; Wernig, M.; Geng, Y.; Moses, A.; Li, C.; Gygi, S. P.; Young, R. A.; Sicinski, P. Cdk1 Controls Global Epigenetic Landscape in Embryonic Stem Cells. Mol Cell 2020, 78 (3), 459–476.e13. 10.1016/j.molcel.2020.03.010.

(46) Chi, Y.; Carter, J. H.; Swanger, J.; Mazin, A. V; Moritz, R. L.; Clurman, B. E. A Novel Landscape of Nuclear Human CDK2 Substrates Revealed by in Situ Phosphorylation. Sci Adv 2020, 6 (16), eaaz9899. 10.1126/sciadv.aaz9899.

(47) Shubbar, E.; Helou, K.; Kovács, A.; Nemes, S.; Hajizadeh, S.; Enerbäck, C.; Einbeigi, Z. High Levels of γ-Glutamyl Hydrolase (GGH) Are Associated with Poor Prognosis and Unfavorable Clinical Outcomes in Invasive Breast Cancer. BMC Cancer 2013, 13, 47. 10.1186/1471-2407-13-47.

(48) Yamauchi, T.; Nio, Y.; Maki, T.; Kobayashi, M.; Takazawa, T.; Iwabu, M.; Okada-Iwabu, M.; Kawamoto, S.; Kubota, N.; Kubota, T.; Ito, Y.; Kamon, J.; Tsuchida, A.; Kumagai, K.; Kozono, H.; Hada, Y.; Ogata, H.; Tokuyama, K.; Tsunoda, M.; Ide, T.; Murakami, K.; Awazawa, M.; Takamoto, I.; Froguel, P.; Hara, K.; Tobe, K.; Nagai, R.; Ueki, K.; Kadowaki, T. Targeted Disruption of AdipoR1 and AdipoR2 Causes Abrogation of Adiponectin Binding and Metabolic Actions. Nat Med 2007, 13 (3), 332–339. 10.1038/nm1557.

(49) Blankestijn, M.; Bloks, V. W.; Struik, D.; Huijkman, N.; Kloosterhuis, N.; Wolters, J. C.; Wanders, R. J. A.; Vaz, F. M.; Islinger, M.; Kuipers, F.; van de Sluis, B.; Groen, A. K.; Verkade, H. J.; Jonker, J. W. Mice with a Deficiency in Peroxisomal Membrane Protein 4 (PXMP4) Display Mild Changes in Hepatic Lipid Metabolism. Sci Rep 2022, 12 (1), 2512. 10.1038/s41598-022-06479-y.

(50) Bustin, M. Chromatin Unfolding and Activation by HMGN(*) Chromosomal Proteins. Trends Biochem Sci 2001, 26 (7), 431–437. 10.1016/s0968-0004(01)01855-2.

(51) Yin, L.; Li, Q.; Mrdenovic, S.; Chu, G. C.-Y.; Wu, B. J.; Bu, H.; Duan, P.; Kim, J.; You, S.; Lewis, M. S.; Liang, G.; Wang, R.; Zhau, H. E.; Chung, L. W. K. KRT13 Promotes Stemness and Drives Metastasis in Breast Cancer through a Plakoglobin/c-Myc Signaling Pathway. Breast Cancer Res 2022, 24 (1), 7. 10.1186/s13058-022-01502-6.

(52) Okita, Y.; Kimura, M.; Xie, R.; Chen, C.; Shen, L. T.-W.; Kojima, Y.; Suzuki, H.; Muratani, M.; Saitoh, M.; Semba, K.; Heldin, C.-H.; Kato, M. The Transcription Factor MAFK Induces EMT and Malignant Progression of Triple-Negative Breast Cancer Cells through Its Target GPNMB. Sci Signal 2017, 10 (474). 10.1126/scisignal.aak9397.

(53) Baniwal, S. K.; Chimge, N.-O.; Jordan, V. C.; Tripathy, D.; Frenkel, B. Prolactin-Induced Protein (PIP) Regulates Proliferation of Luminal A Type Breast Cancer Cells in an Estrogen-Independent Manner. PLoS One 2014, 8 (6), e62361. 10.1371/journal.pone.0062361.

(54) Hwang, S.; Park, S.; Kim, J. H.; Bang, S.-B.; Kim, H.-J.; Ka, N.-L.; Ko, Y.; Kim, S.-S.; Lim, G. Y.; Lee, S.; Shin, Y. K.; Park, S. Y.; Kim, S.; Lee, M.-O. Targeting HMG-CoA Synthase 2 Suppresses Tamoxifen-Resistant Breast Cancer Growth by Augmenting Mitochondrial Oxidative Stress-Mediated Cell Death. Life Sci 2023, 328, 121827. 10.1016/j.lfs.2023.121827.

(55) Ceccaldi, R.; Sarangi, P.; D’Andrea, A. D. The Fanconi Anaemia Pathway: New Players and New Functions. Nat Rev Mol Cell Biol 2016, 17 (6), 337–349. 10.1038/nrm.2016.48.

(56) Mahdessian, D.; Cesnik, A. J.; Gnann, C.; Danielsson, F.; Stenström, L.; Arif, M.; Zhang, C.; Le, T.; Johansson, F.; Schutten, R.; Bäckström, A.; Axelsson, U.; Thul, P.; Cho, N. H.; Carja, O.; Uhlén, M.; Mardinoglu, A.; Stadler, C.; Lindskog, C.; Ayoglu, B.; Leonetti, M. D.; Pontén, F.; Sullivan, D. P.; Lundberg, E. Spatiotemporal Dissection of the Cell Cycle with Single-Cell Proteogenomics. Nature 2021, 590 (7847), 649–654. 10.1038/s41586-021-03232-9.

(57) Ly, T.; Ahmad, Y.; Shlien, A.; Soroka, D.; Mills, A.; Emanuele, M. J.; Stratton, M. R.; Lamond, A. I. A Proteomic Chronology of Gene Expression through the Cell Cycle in Human Myeloid Leukemia Cells. Elife 2014, 3, e01630. 10.7554/eLife.01630.

(58) Santiappillai, N. T.; Abuhammad, S.; Slater, A.; Kirby, L.; McArthur, G. A.; Sheppard, K. E.; Smith, L. K. CDK4/6 Inhibition Reprograms Mitochondrial Metabolism in BRAFV600 Melanoma via a P53 Dependent Pathway. Cancers (Basel) 2021, 13 (3). 10.3390/cancers13030524.

(59) Eeckhoute, J.; Carroll, J. S.; Geistlinger, T. R.; Torres-Arzayus, M. I.; Brown, M. A Cell-Type-Specific Transcriptional Network Required for Estrogen Regulation of Cyclin D1 and Cell Cycle Progression in Breast Cancer. Genes Dev 2006, 20 (18), 2513–2526. 10.1101/gad.1446006.

(60) Mohammadizadeh, F.; Hani, M.; Ranaee, M.; Bagheri, M. Role of Cyclin D1 in Breast Carcinoma. J Res Med Sci 2013, 18 (12), 1021–1025.

(61) Buckley, M. F.; Sweeney, K. J.; Hamilton, J. A.; Sini, R. L.; Manning, D. L.; Nicholson, R. I.; deFazio, A.; Watts, C. K.; Musgrove, E. A.; Sutherland, R. L. Expression and Amplification of Cyclin Genes in Human Breast Cancer. Oncogene 1993, 8 (8), 2127–2133.

(62) Gillett, C.; Fantl, V.; Smith, R.; Fisher, C.; Bartek, J.; Dickson, C.; Barnes, D.; Peters, G. Amplification and Overexpression of Cyclin D1 in Breast Cancer Detected by Immunohistochemical Staining. Cancer Res 1994, 54 (7), 1812–1817.

(63) Tang, J.; Luo, Y.; Long, G.; Zhou, L. MINDY1 Promotes Breast Cancer Cell Proliferation by Stabilizing Estrogen Receptor α. Cell Death Dis 2021, 12 (10), 937. 10.1038/s41419-021-04244-z.

(64) Saleban, M.; Harris, E. L.; Poulter, J. A. D-Type Cyclins in Development and Disease. Genes (Basel) 2023, 14 (7). 10.3390/genes14071445.

(65) Zhang, Q.; Sakamoto, K.; Wagner, K.-U. D-Type Cyclins Are Important Downstream Effectors of Cytokine Signaling That Regulate the Proliferation of Normal and Neoplastic Mammary Epithelial Cells. Mol Cell Endocrinol 2014, 382 (1), 583–592. 10.1016/j.mce.2013.03.016.

(66) Nacht, M.; Ferguson, A. T.; Zhang, W.; Petroziello, J. M.; Cook, B. P.; Gao, Y. H.; Maguire, S.; Riley, D.; Coppola, G.; Landes, G. M.; Madden, S. L.; Sukumar, S. Combining Serial Analysis of Gene Expression and Array Technologies to Identify Genes Differentially Expressed in Breast Cancer. Cancer Res 1999, 59 (21), 5464–5470.

(67) Arafeh, R.; Shibue, T.; Dempster, J. M.; Hahn, W. C.; Vazquez, F. The Present and Future of the Cancer Dependency Map. Nat Rev Cancer 2025, 25 (1), 59–73. 10.1038/s41568-024-00763-x.

(68) Finn, R. S.; Dering, J.; Conklin, D.; Kalous, O.; Cohen, D. J.; Desai, A. J.; Ginther, C.; Atefi, M.; Chen, I.; Fowst, C.; Los, G.; Slamon, D. J. PD 0332991, a Selective Cyclin D Kinase 4/6 Inhibitor, Preferentially Inhibits Proliferation of Luminal Estrogen Receptor-Positive Human Breast Cancer Cell Lines in Vitro. Breast Cancer Res 2009, 11 (5), R77. 10.1186/bcr2419.

(69) Schachter, M. M.; Merrick, K. A.; Larochelle, S.; Hirschi, A.; Zhang, C.; Shokat, K. M.; Rubin, S. M.; Fisher, R. P. A Cdk7-Cdk4 T-Loop Phosphorylation Cascade Promotes G1 Progression. Mol Cell 2013, 50 (2), 250–260. 10.1016/j.molcel.2013.04.003.

(70) Fisher, R. P. Secrets of a Double Agent: CDK7 in Cell-Cycle Control and Transcription. J Cell Sci 2005, 118 (Pt 22), 5171–5180. 10.1242/jcs.02718.

(71) Patel, H.; Abduljabbar, R.; Lai, C.-F.; Periyasamy, M.; Harrod, A.; Gemma, C.; Steel, J. H.; Patel, N.; Busonero, C.; Jerjees, D.; Remenyi, J.; Smith, S.; Gomm, J. J.; Magnani, L.; Győrffy, B.; Jones, L. J.; Fuller-Pace, F.; Shousha, S.; Buluwela, L.; Rakha, E. A.; Ellis, I. O.; Coombes, R. C.; Ali, S. Expression of CDK7, Cyclin H, and MAT1 Is Elevated in Breast Cancer and Is Prognostic in Estrogen Receptor-Positive Breast Cancer. Clin Cancer Res 2016, 22 (23), 5929–5938. 10.1158/1078-0432.CCR-15-1104.

(72) Chen, D.; Riedl, T.; Washbrook, E.; Pace, P. E.; Coombes, R. C.; Egly, J. M.; Ali, S. Activation of Estrogen Receptor Alpha by S118 Phosphorylation Involves a Ligand-Dependent Interaction with TFIIH and Participation of CDK7. Mol Cell 2000, 6 (1), 127–137.

(73) Attia, Y. M.; Shouman, S. A.; Salama, S. A.; Ivan, C.; Elsayed, A. M.; Amero, P.; Rodriguez-Aguayo, C.; Lopez-Berestein, G. Blockade of CDK7 Reverses Endocrine Therapy Resistance in Breast Cancer. Int J Mol Sci 2020, 21 (8). 10.3390/ijms21082974.

(74) Pernas, S.; Hinojo, C.; Pascual, J.; Aksoy, S.; Clack, G.; Mcintosh, S. Fulvestrant with or without the Cyclin-Dependent Kinase 7 (CDK7) Inhibitor Samuraciclib in Advanced Hormone Receptor Positive (HR+) Breast Cancer after CDK4/6 Inhibition: Phase II SUMIT-BC Study; 2025.

(75) Miettinen, T. P.; Peltier, J.; Härtlova, A.; Gierliński, M.; Jansen, V. M.; Trost, M.; Björklund, M. Thermal Proteome Profiling of Breast Cancer Cells Reveals Proteasomal Activation by CDK4/6 Inhibitor Palbociclib. EMBO J 2018, 37 (10). 10.15252/embj.201798359.

(76) Watt, A. C.; Cejas, P.; DeCristo, M. J.; Metzger-Filho, O.; Lam, E. Y. N.; Qiu, X.; BrinJones, H.; Kesten, N.; Coulson, R.; Font-Tello, A.; Lim, K.; Vadhi, R.; Daniels, V. W.; Montero, J.; Taing, L.; Meyer, C. A.; Gilan, O.; Bell, C. C.; Korthauer, K. D.; Giambartolomei, C.; Pasaniuc, B.; Seo, J.-H.; Freedman, M. L.; Ma, C.; Ellis, M. J.; Krop, I.; Winer, E.; Letai, A.; Brown, M.; Dawson, M. A.; Long, H. W.; Zhao, J. J.; Goel, S. CDK4/6 Inhibition Reprograms the Breast Cancer Enhancer Landscape by Stimulating AP-1 Transcriptional Activity. Nat Cancer 2021, 2 (1), 34–48. 10.1038/s43018-020-00135-y.

(77) AbuHammad, S.; Cullinane, C.; Martin, C.; Bacolas, Z.; Ward, T.; Chen, H.; Slater, A.; Ardley, K.; Kirby, L.; Chan, K. T.; Brajanovski, N.; Smith, L. K.; Rao, A. D.; Lelliott, E. J.; Kleinschmidt, M.; Vergara, I. A.; Papenfuss, A. T.; Lau, P.; Ghosh, P.; Haupt, S.; Haupt, Y.; Sanij, E.; Poortinga, G.; Pearson, R. B.; Falk, H.; Curtis, D. J.; Stupple, P.; Devlin, M.; Street, I.; Davies, M. A.; McArthur, G. A.; Sheppard, K. E. Regulation of PRMT5-MDM4 Axis Is Critical in the Response to CDK4/6 Inhibitors in Melanoma. Proc Natl Acad Sci U S A 2019, 116 (36), 17990–18000. 10.1073/pnas.1901323116.

(78) Lin, C.-C.; Chang, T.-C.; Wang, Y.; Guo, L.; Gao, Y.; Bikorimana, E.; Lemoff, A.; Fang, Y. V; Zhang, H.; Zhang, Y.; Ye, D.; Soria-Bretones, I.; Servetto, A.; Lee, K.-M.; Luo, X.; Otto, J. J.; Akamatsu, H.; Napolitano, F.; Mani, R.; Cescon, D. W.; Xu, L.; Xie, Y.; Mendell, J. T.; Hanker, A. B.; Arteaga, C. L. PRMT5 Is an Actionable Therapeutic Target in CDK4/6 Inhibitor-Resistant ER+/RB-Deficient Breast Cancer. Nat Commun 2024, 15 (1), 2287. 10.1038/s41467-024-46495-2.

(79) Guccione, E.; Richard, S. The Regulation, Functions and Clinical Relevance of Arginine Methylation. Nat Rev Mol Cell Biol 2019, 20 (10), 642–657. 10.1038/s41580-019-0155-x.

(80) Yu, S.; Si, Y.; Xu, M.; Wang, Y.; Liu, C.; Bi, C.; Sun, M.; Sun, H. Downregulation of the Splicing Regulator NSRP1 Confers Resistance to CDK4/6 Inhibitors via Activation of Interferon Signaling in Breast Cancer. J Biol Chem 2025, 301 (1), 108070. 10.1016/j.jbc.2024.108070.

(81) Tan, K.; Sebat, J.; Wilkinson, M. F. Cell Type- and Factor-Specific Nonsense-Mediated RNA Decay. Nucleic Acids Res 2025, 53 (9). 10.1093/nar/gkaf395.

(82) Fuentes, N.; Silveyra, P. Estrogen Receptor Signaling Mechanisms. Adv Protein Chem Struct Biol 2019, 116, 135–170. 10.1016/bs.apcsb.2019.01.001.

(83) Järvensivu, P.; Saloniemi-Heinonen, T.; Awosanya, M.; Koskimies, P.; Saarinen, N.; Poutanen, M. HSD17B1 Expression Enhances Estrogen Signaling Stimulated by the Low Active Estrone, Evidenced by an Estrogen Responsive Element-Driven Reporter Gene in Vivo. Chem Biol Interact 2015, 234, 126–134. 10.1016/j.cbi.2015.01.008.

(84) Kannan, N.; Kang, J.; Kong, X.; Tang, J.; Perry, J. K.; Mohankumar, K. M.; Miller, L. D.; Liu, E. T.; Mertani, H. C.; Zhu, T.; Grandison, P. M.; Liu, D.-X.; Lobie, P. E. Trefoil Factor 3 Is Oncogenic and Mediates Anti-Estrogen Resistance in Human Mammary Carcinoma. Neoplasia 2010, 12 (12), 1041–1053. 10.1593/neo.10916.

(85) UniProt Consortium. UniProt: The Universal Protein Knowledgebase in 2025. Nucleic Acids Res 2025, 53 (D1), D609–D617. 10.1093/nar/gkae1010.

(86) Tyanova, S.; Temu, T.; Cox, J. The MaxQuant Computational Platform for Mass Spectrometry-Based Shotgun Proteomics. Nat Protoc 2016, 11 (12), 2301–2319. 10.1038/nprot.2016.136.

(87) Cox, J.; Mann, M. MaxQuant Enables High Peptide Identification Rates, Individualized p.p.b.-Range Mass Accuracies and Proteome-Wide Protein Quantification. Nat Biotechnol 2008, 26 (12), 1367–1372. 10.1038/nbt.1511.

(88) Mudge, J. M.; Carbonell-Sala, S.; Diekhans, M.; Martinez, J. G.; Hunt, T.; Jungreis, I.; Loveland, J. E.; Arnan, C.; Barnes, I.; Bennett, R.; Berry, A.; Bignell, A.; Cerdán-Vélez, D.; Cochran, K.; Cortés, L. T.; Davidson, C.; Donaldson, S.; Dursun, C.; Fatima, R.; Hardy, M.; Hebbar, P.; Hollis, Z.; James, B. T.; Jiang, Y.; Johnson, R.; Kaur, G.; Kay, M.; Mangan, R. J.; Maquedano, M.; Gómez, L. M.; Mathlouthi, N.; Merritt, R.; Ni, P.; Palumbo, E.; Perteghella, T.; Pozo, F.; Raj, S.; Sisu, C.; Steed, E.; Sumathipala, D.; Suner, M.-M.; Uszczynska-Ratajczak, B.; Wass, E.; Yang, Y. T.; Zhang, D.; Finn, R. D.; Gerstein, M.; Guigó, R.; Hubbard, T. J. P.; Kellis, M.; Kundaje, A.; Paten, B.; Tress, M. L.; Birney, E.; Martin, F. J.; Frankish, A. GENCODE 2025: Reference Gene Annotation for Human and Mouse. Nucleic Acids Res 2025, 53 (D1), D966–D975. 10.1093/nar/gkae1078.

(89) Chen, S.; Zhou, Y.; Chen, Y.; Gu, J. Fastp: An Ultra-Fast All-in-One FASTQ Preprocessor. Bioinformatics 2018, 34 (17), i884–i890. 10.1093/bioinformatics/bty560.

(90) Dobin, A.; Davis, C. A.; Schlesinger, F.; Drenkow, J.; Zaleski, C.; Jha, S.; Batut, P.; Chaisson, M.; Gingeras, T. R. STAR: Ultrafast Universal RNA-Seq Aligner. Bioinformatics 2013, 29 (1), 15–21. 10.1093/bioinformatics/bts635.

(91) Li, H.; Handsaker, B.; Wysoker, A.; Fennell, T.; Ruan, J.; Homer, N.; Marth, G.; Abecasis, G.; Durbin, R.; 1000 Genome Project Data Processing Subgroup. The Sequence Alignment/Map Format and SAMtools. Bioinformatics 2009, 25 (16), 2078–2079. 10.1093/bioinformatics/btp352.

(92) Ewels, P.; Magnusson, M.; Lundin, S.; Käller, M. MultiQC: Summarize Analysis Results for Multiple Tools and Samples in a Single Report. Bioinformatics 2016, 32 (19), 3047–3048. 10.1093/bioinformatics/btw354.

(93) Patro, R.; Duggal, G.; Love, M. I.; Irizarry, R. A.; Kingsford, C. Salmon Provides Fast and Bias-Aware Quantification of Transcript Expression. Nat Methods 2017, 14 (4), 417–419. 10.1038/nmeth.4197.

(94) R Core Team. R: A Language and Environment for Statistical Computing. Vienna 2023.

(95) Love, M. I.; Huber, W.; Anders, S. Moderated Estimation of Fold Change and Dispersion for RNA-Seq Data with DESeq2. Genome Biol 2014, 15 (12), 550. 10.1186/s13059-014-0550-8.

(96) Buenrostro, J. D.; Giresi, P. G.; Zaba, L. C.; Chang, H. Y.; Greenleaf, W. J. Transposition of Native Chromatin for Fast and Sensitive Epigenomic Profiling of Open Chromatin, DNA-Binding Proteins and Nucleosome Position. Nat Methods 2013, 10 (12), 1213–1218. 10.1038/nmeth.2688.

(97) Corces, M. R.; Trevino, A. E.; Hamilton, E. G.; Greenside, P. G.; Sinnott-Armstrong, N. A.; Vesuna, S.; Satpathy, A. T.; Rubin, A. J.; Montine, K. S.; Wu, B.; Kathiria, A.; Cho, S. W.; Mumbach, M. R.; Carter, A. C.; Kasowski, M.; Orloff, L. A.; Risca, V. I.; Kundaje, A.; Khavari, P. A.; Montine, T. J.; Greenleaf, W. J.; Chang, H. Y. An Improved ATAC-Seq Protocol Reduces Background and Enables Interrogation of Frozen Tissues. Nat Methods 2017, 14 (10), 959–962. 10.1038/nmeth.4396.

(98) Andrews, S. FastQC: A Quality Control Tool for High Throughput Sequence Data.

(99) Langmead, B.; Salzberg, S. L. Fast Gapped-Read Alignment with Bowtie 2. Nat Methods 2012, 9 (4), 357–359. 10.1038/nmeth.1923.

(100) Li, H.; Handsaker, B.; Wysoker, A.; Fennell, T.; Ruan, J.; Homer, N.; Marth, G.; Abecasis, G.; Durbin, R.; 1000 Genome Project Data Processing Subgroup. The Sequence Alignment/Map Format and SAMtools. Bioinformatics 2009, 25 (16), 2078–2079. 10.1093/bioinformatics/btp352.

(101) Broad Institute. Picard Toolkit.

(102) Ramírez, F.; Dündar, F.; Diehl, S.; Grüning, B. A.; Manke, T. DeepTools: A Flexible Platform for Exploring Deep-Sequencing Data. Nucleic Acids Res 2014, 42 (Web Server issue), W187–91. 10.1093/nar/gku365.

(103) Zhang, Y.; Liu, T.; Meyer, C. A.; Eeckhoute, J.; Johnson, D. S.; Bernstein, B. E.; Nusbaum, C.; Myers, R. M.; Brown, M.; Li, W.; Liu, X. S. Model-Based Analysis of ChIP-Seq (MACS). Genome Biol 2008, 9 (9), R137. 10.1186/gb-2008-9-9-r137.

(104) Quinlan, A. R.; Hall, I. M. BEDTools: A Flexible Suite of Utilities for Comparing Genomic Features. Bioinformatics 2010, 26 (6), 841–842. 10.1093/bioinformatics/btq033.

(105) Liao, Y.; Smyth, G. K.; Shi, W. The Subread Aligner: Fast, Accurate and Scalable Read Mapping by Seed-and-Vote. Nucleic Acids Res 2013, 41 (10), e108. 10.1093/nar/gkt214.

(106) Liao, Y.; Smyth, G. K.; Shi, W. FeatureCounts: An Efficient General Purpose Program for Assigning Sequence Reads to Genomic Features. Bioinformatics 2014, 30 (7), 923–930. 10.1093/bioinformatics/btt656.

(107) Yu, G.; Wang, L.-G.; Han, Y.; He, Q.-Y. ClusterProfiler: An R Package for Comparing Biological Themes among Gene Clusters. OMICS 2012, 16 (5), 284–287. 10.1089/omi.2011.0118.

(108) Yu, G.; Wang, L.-G.; Han, Y.; He, Q.-Y. ClusterProfiler: An R Package for Comparing Biological Themes among Gene Clusters. OMICS 2012, 16 (5), 284–287. 10.1089/omi.2011.0118.

(109) Korotkevich, G.; Sukhov, V.; Sergushichev, A. Fast Gene Set Enrichment Analysis. bioRxiv 2019.

(110) Liberzon, A.; Subramanian, A.; Pinchback, R.; Thorvaldsdóttir, H.; Tamayo, P.; Mesirov, J. P. Molecular Signatures Database (MSigDB) 3.0. Bioinformatics 2011, 27 (12), 1739–1740. 10.1093/bioinformatics/btr260.

(111) Brennan, P. DrawProteins: A Bioconductor/R Package for Reproducible and Programmatic Generation of Protein Schematics. F1000Res 2018, 7, 1105. 10.12688/f1000research.14541.1.

(112) Lane, K. R.; Yu, Y.; Lackey, P. E.; Chen, X.; Marzluff, W. F.; Cook, J. G. Cell Cycle-Regulated Protein Abundance Changes in Synchronously Proliferating HeLa Cells Include Regulation of Pre-MRNA Splicing Proteins. PLoS One 2013, 8 (3), e58456. 10.1371/journal.pone.0058456.

(113) DepMap; Broad. DepMap 24Q4 Public. Figshare+. Dataset.

(114) Perez, G.; Barber, G. P.; Benet-Pages, A.; Casper, J.; Clawson, H.; Diekhans, M.; Fischer, C.; Gonzalez, J. N.; Hinrichs, A. S.; Lee, C. M.; Nassar, L. R.; Raney, B. J.; Speir, M. L.; van Baren, M. J.; Vaske, C. J.; Haussler, D.; Kent, W. J.; Haeussler, M. The UCSC Genome Browser Database: 2025 Update. Nucleic Acids Res 2025, 53 (D1), D1243–D1249. 10.1093/nar/gkae974.

(115) Untergasser, A.; Cutcutache, I.; Koressaar, T.; Ye, J.; Faircloth, B. C.; Remm, M.; Rozen, S. G. Primer3--New Capabilities and Interfaces. Nucleic Acids Res 2012, 40 (15), e115. 10.1093/nar/gks596.

