## Supplemental Figures, Legends, and Table Legends for "Charting the Multi-level Molecular Response to Palbociclib in ER-Positive Breast Cancer"

### Supplementary Materials

Archishma Kavalipati[1,2*], Amy Aponte[2,*], Michael E. Sullivan[2], Sarah L. Whittington[3], José C. Martínez[3,4], Grant Goda[4], Maria M. Aleman[2], Michael J. Emanuele[2,3,#], Daniel Dominguez[1,2,3,6,#]

**Supplementary Tables**

**Table S1:** Differential gene expression after Palbo treatment.

**Table S2:** Differential chromatin accessibility after Palbo treatment.

**Table S3:** Alternative splicing after Palbo treatment.

**Table S4:** Differential proteomic expression after Palbo treatment.

**Table S5:** Reagents, primers, and software table. Tabs are in order as follows: antibodies, chemicals, recombinant proteins, commercial assays/reagents, primers for splicing analysis, R software/packages used.

**Supplementary Figure Legends**

**Figure S1**

**A.** Upper: PCA plot of RNA-seq data. Lower: PCA plot of ATAC-seq data.

**B.** Top Gene Ontology (GO) terms enriched in downregulated genes compared to all genes detected. The p-value and q-value cutoffs were both set to 0.05 and the ontology "BP" (biological process) was used. Top 5 enriched terms are shown.

**C.** Left: Overlap between downregulated genes and periodic genes with peak expression in G1/S phase (p = 1.21e-24, hypergeometric test). Right: Overlap between downregulated genes and periodic genes with peak expression in G2/M phase (p = 3.53e-34, hypergeometric test).

**D.** UpSet plot showing overlaps between Palbo-downregulated genes and periodic genes from external datasets: Whitfield et al, 2002; Dominguez et al, 2016; Boström et al, 2017; and Li, Wang and Wang et al, 2022. The 18 genes overlapping all 5 datasets are highlighted.

**E.** ATAC-seq volcano plot showing differential accessibility in chromatin regions after Palbo treatment. Genes overlapping open chromatin regions are labeled if significant at mRNA level and abs(log2FoldChange) of chromatin region $\geq 0.6$.

**F.** Scatter plot of accessibility (ATAC-seq) and mRNA level log2FoldChange (Pearson correlation = 0.26, p = 2.2e-16)

**G:** Heatmap showing pyPAGE inference of transcription factors (TFs) involved in regulation of differentially expressed genes. Top annotation: Average log2FoldChange of the genes in each bin. The top 10 inferred TFs are shown.

**Figure S2**

**A.** Ratios of significant events to all events detected using rMATS.

**B.** Overlap between periodically spliced genes from Dominguez et al. 2016 and Palbo-induced alternatively spliced genes. p = 3.34e-04, hypergeometric test.

**C.** Overlap between periodic splicing events identified from Dominguez et al. 2016 and Palbo-induced alternative splicing events. p = 0.24, hypergeometric test.

**D.** Left: visualization of genome coverage in ECT2 exon skipping event. Right: validation of alternative splicing by quantitative RT-PCR and corresponding quantification (p = 1.01e-02, Student's two-sided t-test).

**E.** Upper: genome coverage of 3 AURKA splicing events overlapping similar genomic intervals. Lower; validation of AURKA exon skipping RT-PCR and corresponding quantification (p=1.19e-04, Student's two-sided t-test). Note that isoform level resolution was not possible due to similarity of size for inclusion products.

**Figure S3**

**A:** Western blot showing of indicated proteins indicating cycle arrest after Palbo treatment from samples used for MS.

**B:** Subset of correlation plot from **Fig. 3C** showing genes with decreased mRNA and increased protein abundance.

**C:** Subset of correlation plot from **Fig. 3C** showing genes with increased mRNA and decreased protein abundance.

**D:** UpSet plot showing intersections between downregulated proteins and periodic proteins identified in external datasets: Ly et al. 2014, Lane et al. 2013, Mahdessian et al. 2021. 9 proteins overlapping all datasets are highlighted. Intersections with a degree ≥ 3 are shown.

**E:** UpSet plot showing intersections between upregulated proteins and cycling proteins identified in external datasets (described above). 2 proteins identified that overlap with Palbo-upregulated proteins are shown. Intersections with a degree ≥ 2 are shown.

**Figure S4**

**A:** p-adjusted values (KS test, Benjamini-Hochberg correction) obtained after 500 trials comparing true Palbo-downregulated genes at mRNA level to a null distribution obtained from matching to genes with similar average FPKMs to the DMSO condition. Red line: p = 5.89e-11, KS test, downregulated genes compared to all non-downregulated genes.

**B:** p-adjusted values (KS test, Benjamini-Hochberg correction) obtained after 500 trials comparing true Palbo-upregulated genes at mRNA level to a null distribution obtained from matching to genes with similar average FPKMs to the DMSO condition. Red line: p = 4.06e-04, KS test, downregulated genes compared to all non-downregulated genes.

**C:** p-adjusted values (KS test, Benjamini-Hochberg correction) obtained after 500 trials comparing true Palbo-downregulated genes at protein level to a null distribution obtained from matching to genes with similar average abundances to the DMSO condition. Red line: p = 1.72e-07, KS test, downregulated genes compared to all non-downregulated genes.

**D:** p-adjusted values (KS test, Benjamini-Hochberg correction) obtained after 500 trials comparing true Palbo-upregulated genes at protein level to a null distribution obtained from matching to genes with similar average abundances to the DMSO condition. Red line: p = 1.6e-02, KS test, downregulated genes compared to all non-downregulated genes.

#### Figure S1

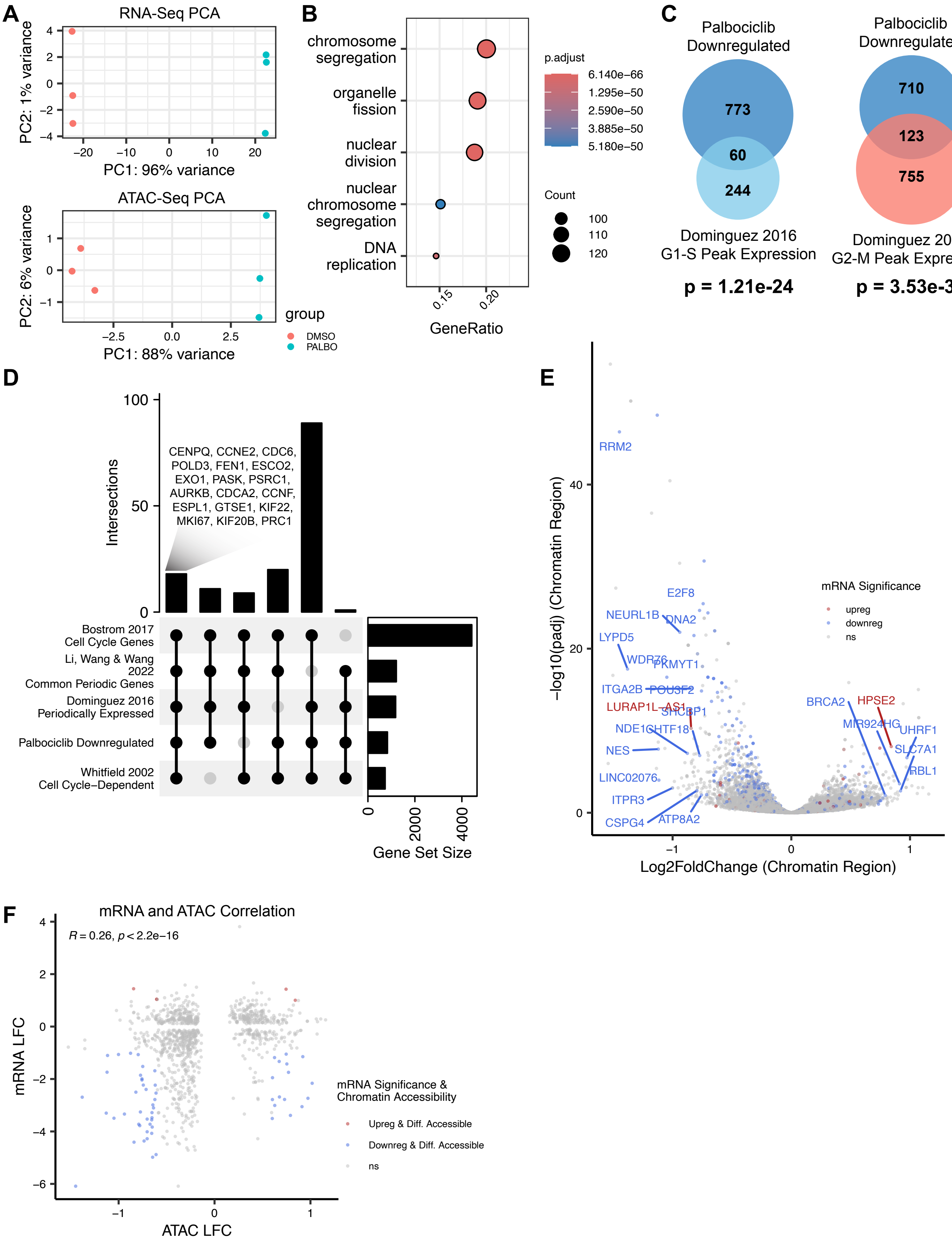

A
RNA-Seq PCA
PC1: 96% variance
PC2: 1% variance
ATAC-Seq PCA
PC1: 88% variance
PC2: 6% variance
group
DMSO
PALBO
B
chromosome segregation
organelle fission
nuclear division
nuclear chromosome segregation
DNA replication
GeneRatio
p.adjust
6.140e−66
1.295e−50
2.590e−50
3.885e−50
5.180e−50
Count
100
110
120
C
Palbociclib Downregulated
773
60
244
Dominguez 2016 G1-S Peak Expression
p = 1.21e-24
Palbociclib Downregulated
710
123
755
Dominguez 2016 G2-M Peak Expression
p = 3.53e-34
D
Intersections
CENPQ, CCNE2, CDC6, POLD3, FEN1, ESCO2, EXO1, PASK, PSRC1, AURKB, CDCA2, CCNF, ESPL1, GTSE1, KIF22, MKI67, KIF20B, PRC1
Bostrom 2017 Cell Cycle Genes
Li, Wang & Wang 2022 Common Periodic Genes
Dominguez 2016 Periodically Expressed
Palbociclib Downregulated
Whitfield 2002 Cell Cycle−Dependent
Gene Set Size
E
−log10(padj) (Chromatin Region)
Log2FoldChange (Chromatin Region)
mRNA Significance
upreg
downreg
ns
RRM2
E2F8
NEURL1B
DNA2
LYPD5
WDR76
PKMYT1
ITGA2B
POU3F2
LURAP1L−AS1
SHCBP1
NDE1
GHTF18
NES
LINC02076
ITPR3
CSPG4
ATP8A2
BRCA2
HPSE2
MIR924HG
UHRF1
SLC7A1
RBL1
F
mRNA and ATAC Correlation
R = 0.26, p < 2.2e−16
mRNA LFC
ATAC LFC
mRNA Significance & Chromatin Accessibility
Upreg & Diff. Accessible
Downreg & Diff. Accessible
ns

#### Figure S2

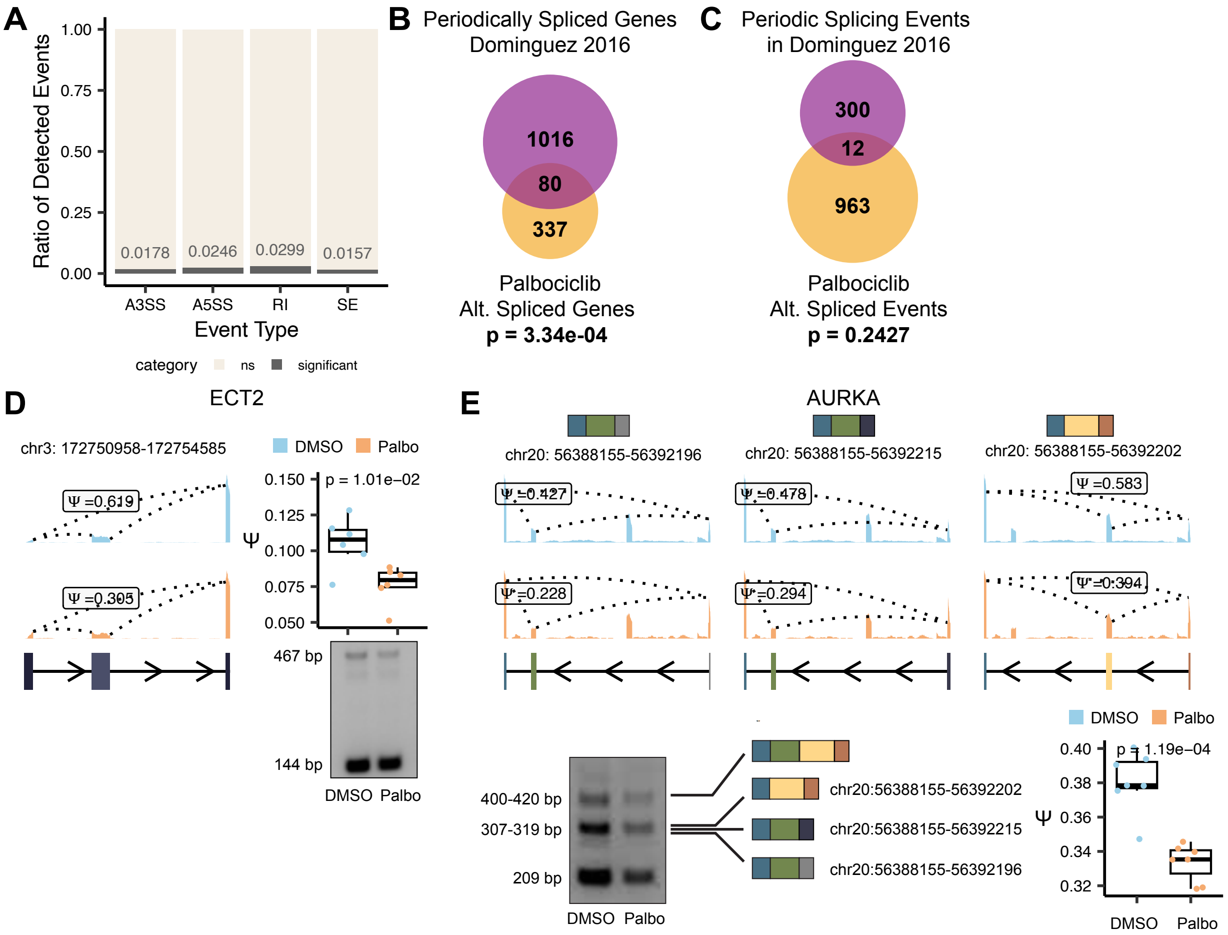

A
Ratio of Detected Events
1.00
0.75
0.50
0.25
0.00
0.0178
0.0246
0.0299
0.0157
A3SS
A5SS
RI
SE
Event Type
category
ns
significant
B
Periodically Spliced Genes
Dominguez 2016
1016
80
337
Palbociclib
Alt. Spliced Genes
p = 3.34e-04
C
Periodic Splicing Events
in Dominguez 2016
300
12
963
Palbociclib
Alt. Spliced Events
p = 0.2427
D
ECT2
chr3: 172750958-172754585
DMSO
Palbo
Ψ =0.619
Ψ =0.305
p = 1.01e−02
0.150
0.125
0.100
0.075
0.050
Ψ
467 bp
144 bp
DMSO
Palbo
E
AURKA
chr20: 56388155-56392196
chr20: 56388155-56392215
chr20: 56388155-56392202
Ψ =0.427
Ψ =0.478
Ψ =0.583
Ψ =0.228
Ψ =0.294
Ψ =0.394
400-420 bp
307-319 bp
209 bp
DMSO
Palbo
chr20:56388155-56392202
chr20:56388155-56392215
chr20:56388155-56392196
p = 1.19e−04
0.40
0.38
0.36
0.34
0.32
Ψ

**Figure S3**

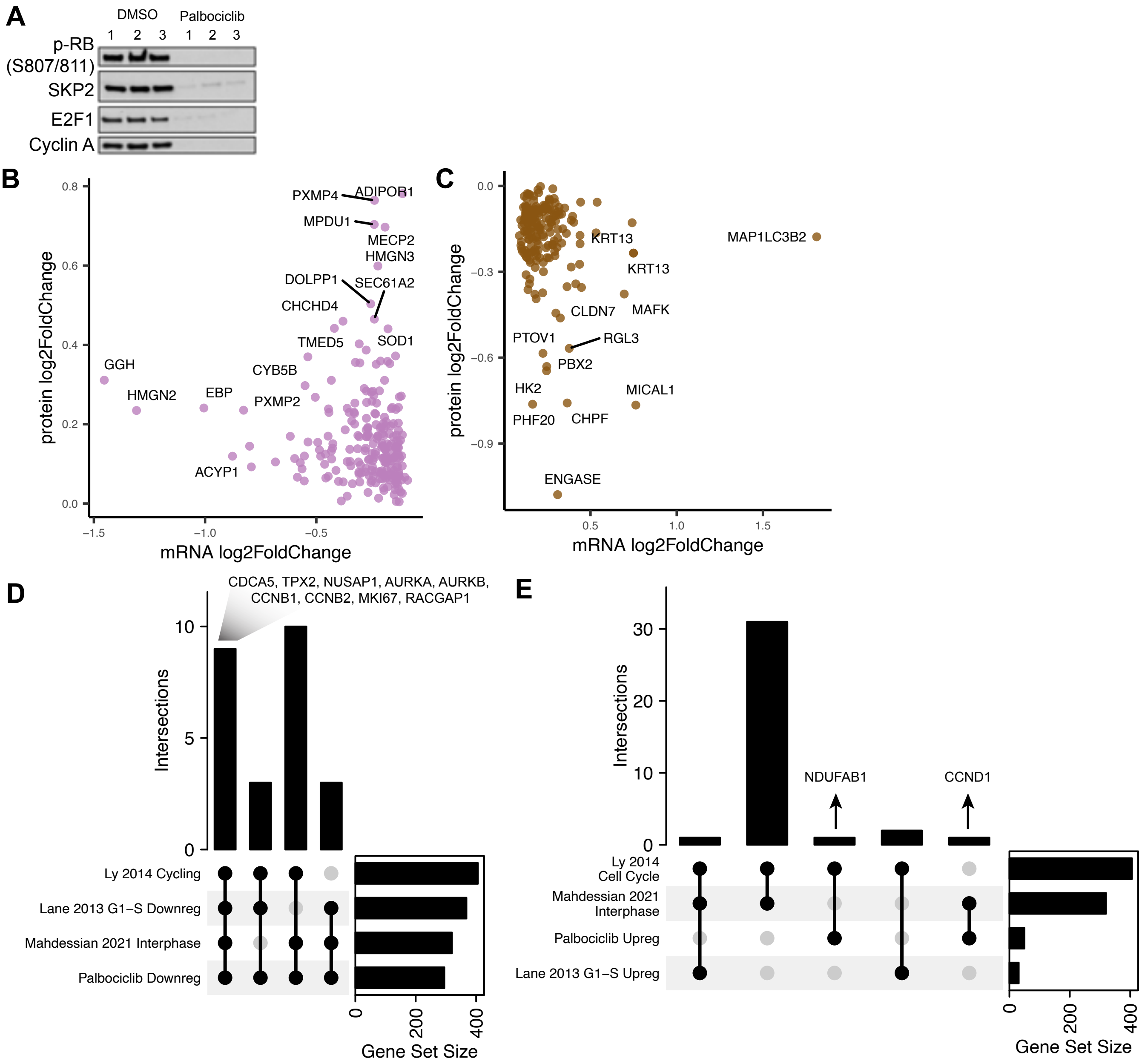
A
DMSO
Palbociclib
1 2 3 1 2 3
p-RB (S807/811)
SKP2
E2F1
Cyclin A
B
protein log2FoldChange
mRNA log2FoldChange
PXMP4
ADIPOR1
MPDU1
MECP2
HMGN3
DOLPP1
SEC61A2
CHCHD4
TMED5
SOD1
GGH
CYB5B
HMGN2
EBP
PXMP2
ACYP1
C
protein log2FoldChange
mRNA log2FoldChange
KRT13
MAP1LC3B2
KRT13
CLDN7
MAFK
PTOV1
RGL3
PBX2
HK2
MICAL1
PHF20
CHPF
ENGASE
D
CDCA5, TPX2, NUSAP1, AURKA, AURKB, CCNB1, CCNB2, MKI67, RACGAP1
Intersections
Ly 2014 Cycling
Lane 2013 G1−S Downreg
Mahdessian 2021 Interphase
Palbociclib Downreg
Gene Set Size
E
Intersections
NDUFAB1
CCND1
Ly 2014 Cell Cycle
Mahdessian 2021 Interphase
Palbociclib Upreg
Lane 2013 G1−S Upreg
Gene Set Size

### Figure S4

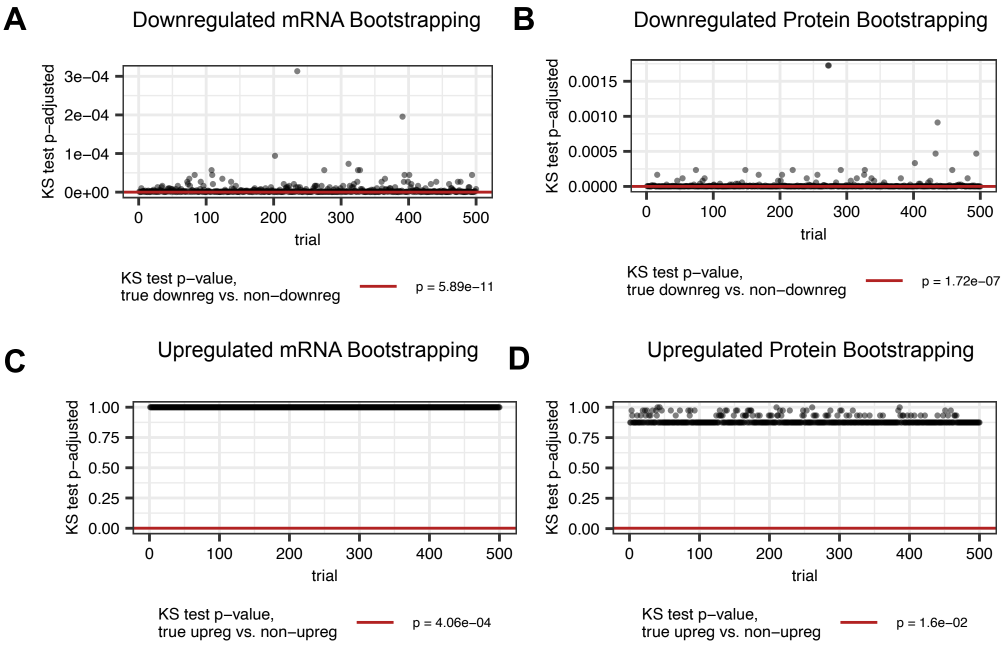
A
Downregulated mRNA Bootstrapping
KS test p−adjusted
3e−04
2e−04
1e−04
0e+00
0
100
200
300
400
500
trial
KS test p−value,
true downreg vs. non−downreg
p = 5.89e−11
B
Downregulated Protein Bootstrapping
KS test p−adjusted
0.0015
0.0010
0.0005
0.0000
0
100
200
300
400
500
trial
KS test p−value,
true downreg vs. non−downreg
p = 1.72e−07
C
Upregulated mRNA Bootstrapping
KS test p−adjusted
1.00
0.75
0.50
0.25
0.00
0
100
200
300
400
500
trial
KS test p−value,
true upreg vs. non−upreg
p = 4.06e−04
D
Upregulated Protein Bootstrapping
KS test p−adjusted
1.00
0.75
0.50
0.25
0.00
0
100
200
300
400
500
trial
KS test p−value,
true upreg vs. non−upreg
p = 1.6e−02
